# Targeting MTHFD2 disrupts mitochondrial redox homeostasis and restores venetoclax sensitivity in acute myeloid leukemia

**DOI:** 10.64898/2026.03.18.712743

**Authors:** Judith O. Sokei, Orsola di Martino, Mary Basse, Nishanth Gabriel, Liana Valin, Conner R. York, Nancy B.J. Arthur, Wujuan Zhang, Aaron R. Goldman, Francesca Ferraro, Stephen M. Sykes

## Abstract

One-carbon metabolism is frequently dysregulated in human cancer including acute myeloid leukemia. However, the mitochondrial mechanisms by which one-carbon enzymes support leukemia survival and therapeutic response remain incompletely defined. Here, we report that the one-carbon metabolism enzyme MTHFD2 is a critical regulator of acute myeloid leukemia nucleotide metabolism, redox homeostasis, and disease progression. We show that genetic ablation of MTHFD2 suppresses acute myeloid leukemia cell proliferation *in vitro* and significantly delays leukemia onset in a genetically engineered mouse model, while sparing healthy hematopoietic stem and progenitor cell function. Stable isotope tracing demonstrates that MTHFD2 supports *de novo* purine synthesis and sustains mitochondrial NADH and NADPH production. Consistent with this role, MTHFD2 inhibition increases mitochondrial superoxide levels, and combined purine supplementation and mitochondrial reactive oxygen species neutralization rescues acute myeloid leukemia cell viability. We also demonstrate that the small-molecule inhibitor DS18561882 directly inhibits mitochondrial MTHFD2 activity and phenocopies genetic deletion. DS18561882 exhibits activity across a cohort of 60 primary AML patient samples, synergizes with venetoclax in treatment-naïve acute myeloid leukemia, and restores venetoclax sensitivity in resistant AML models. These findings establish mitochondrial MTHFD2 as a genetically validated, therapeutically targetable metabolic vulnerability in acute myeloid leukemia and support targeting mitochondrial one-carbon metabolism to enhance and restore venetoclax response.

## Introduction

The identification and subsequent targeting of metabolic vulnerabilities has emerged as a focal point in cancer biology^1–3^. While many aspects of cellular metabolism are altered in human cancer, one-carbon metabolism is frequently co-opted in many tumor settings^4^. This multifaceted metabolic network maintains intracellular pools of deoxythymidine triphosphate (dTTP), *de novo* purine synthesis, redox homeostasis, and epigenetic regulation^4–6^. Several cancers, including acute myeloid leukemia (AML), and those of breast, colon, esophageal, and lung, rely on one-carbon metabolism to synthesize purines and dTTP to support high rates of DNA synthesis^7–12^. Certain cancers rely on one-carbon metabolism to generate NADH and NADPH to support additional metabolic processes and maintain redox homeostasis^9,12–15^. The central backbone of one-carbon metabolism is comprised of two parallel pathways: a mitochondria arm and a cytosolic arm. Although many of the same metabolic intermediates can be generated by both pathways, many cancer cells catabolize serine through the mitochondrial arm^9,10^. Consistent with this observation, two key mitochondrial one-carbon metabolic enzymes, MTHFD2 and SHMT2, are frequently over-expressed in several human cancers including AML^14,16,17^.

AML remains the deadliest form of leukemia in adults and children^18–20^. We and others have reported that certain aggressive subtypes of AML, including those bearing KMT2A-rearrangements or internal tandem duplications of the *FLT3* gene (FLT3^ITD^), are highly dependent on one-carbon metabolism as well as MTHFD2^11,21^. Bonagas et al. recently developed a chemical compound, TH9619, that inhibits MTHFD2 and displays anti-leukemia activity in preclinical models of AML^22^. Mechanistically, they showed that TH9619 causes formate accumulation and thymidine pool depletion leading to DNA replication fork stalling and subsequent AML cell cycle arrest. However, follow-up analyses revealed that TH9619 effectively blocks nuclear MTHFD2 and cytosolic MTHFD1 by cannot enter the mitochondria^23^. As a result, the mitochondrial role of MTHFD2 in AML remains unresolved.

To define the role of MTHFD2 in AML cell biology, we employed a genetic approach to assess the impact of MTHFD2 deletion on normal and malignant hematopoiesis. We report that MTHFD2 ablation does not impact healthy murine hematopoiesis but significantly impedes disease progression in a genetically engineered mouse model of AML. We also show that loss of MTHFD2 blocks purine synthesis and diminishes mitochondrial NADH and NADPH generation, thereby disrupting local redox homeostasis. Confirming that AML cells rely on MTHFD2 to maintain purine pools and mitochondrial redox balance, we show that combining exogenous purine supplementation with mitochondrial reactive oxygen species (ROS) neutralization mitigates the anti-leukemia effects of MTHFD2 inhibition. We also demonstrate that the MTHFD2 inhibitor DS18561882, is able to penetrate the mitochondria, displays anti-leukemia activity as a single agent, and cooperates with the FDA-approved anti-AML therapy Venetoclax reducing patient-derived AML (PD-AML) cell viability. Furthermore, we show that AML cells that have developed resistance to Venetoclax are highly sensitive to DS18561882. Collectively, these studies provide new mechanistic insights into the role of MTHFD2 in AML and establish mitochondrial MTHFD2 as a therapeutically vulnerability in AML.

## Results

### MTHFD2 supports AML

We first employed a CRISPR/Cas9 approach to assess how loss of *MTHFD2* impacts AML cell biology. Briefly, the human AML cell lines MOLM14, NOMO1, OCI-AML2 and OCI-AML3 were engineered to express Cas9 under the control of a doxycycline (DOX)-inducible promoter. Stable Cas9-expressing cell lines were then transduced with lentiviral constructs co-expressing GFP and either control (sgNT) or MTHFD2-targeting (sgMTHFD2-1 or sgMTHFD2-6) guide RNAs (sgRNAs). Total MTHFD2 protein levels, including mitochondrial MTHFD2, were effectively reduced by both MTHFD2-targeting sgRNAs (Figure 1A and 1B and Supplementary Figure 1A-1C). Deletion of *MTHFD2* impaired the expansion of multiple AML cell lines, including MOLM14, NOMO1, OCI-AML2 and OCI-AML3 (Figure 1A and 1B and Supplementary Figure 1B and 1C). This reduction in cell expansion was associated with reduced cell cycling and increased cell death, as measured by BrdU incorporation and Annexin V staining, respectively (Figure 1C and 1D and Supplementary Figure 1D and 1E). To assess the impact of *MTHFD2* deletion on AML biology *in vivo*, we obtained mice carrying loxP sites flanking exon 2 of *Mthfd2*^24^ and further engineered them to express the Mx1-Cre transgene^25^. We used recombinant retroviruses expressing MLL-AF9^26^ to transform bone marrow-derived hematopoietic progenitors from *Mthfd2^+/+^*;Mx1-Cre and *Mthfd2^flox/flox^*;Mx1-Cre mice. Frank leukemia cells from both conditions were separately transplanted into sub-lethally irradiated syngeneic recipient mice. At 10 days post-transplant, mice from each group were administered poly-inosine-poly-cytidine (pIpC) to induce Cre-driven *Mthfd2* gene deletion. Compared to controls (*Mthfd2^+/+^*;MA9), deletion of *Mthfd2* (*Mthfd2^Δ/Δ^*;MA9) resulted in an overall delay in the onset of disease (Figure 1E) and significantly lower white blood cell counts (Figure 1F). PCR-based excision analysis of leukemia cells recovered from the bone marrow (BM) of four *Mthfd2^+/+^*;MA9 and four *Mthfd2^Δ/Δ^*;MA9 leukemic mice showed that *Mthfd2* was predominantly deleted in all four *Mthfd2^Δ/Δ^*;MA9 mice, although two mice displayed a visible portion of unexcised alleles (Supplementary Figure 1F). These findings demonstrate that MTHFD2 functionally supports leukemic cell proliferation, survival, and disease progression in these AML models.

**Figure 1.**
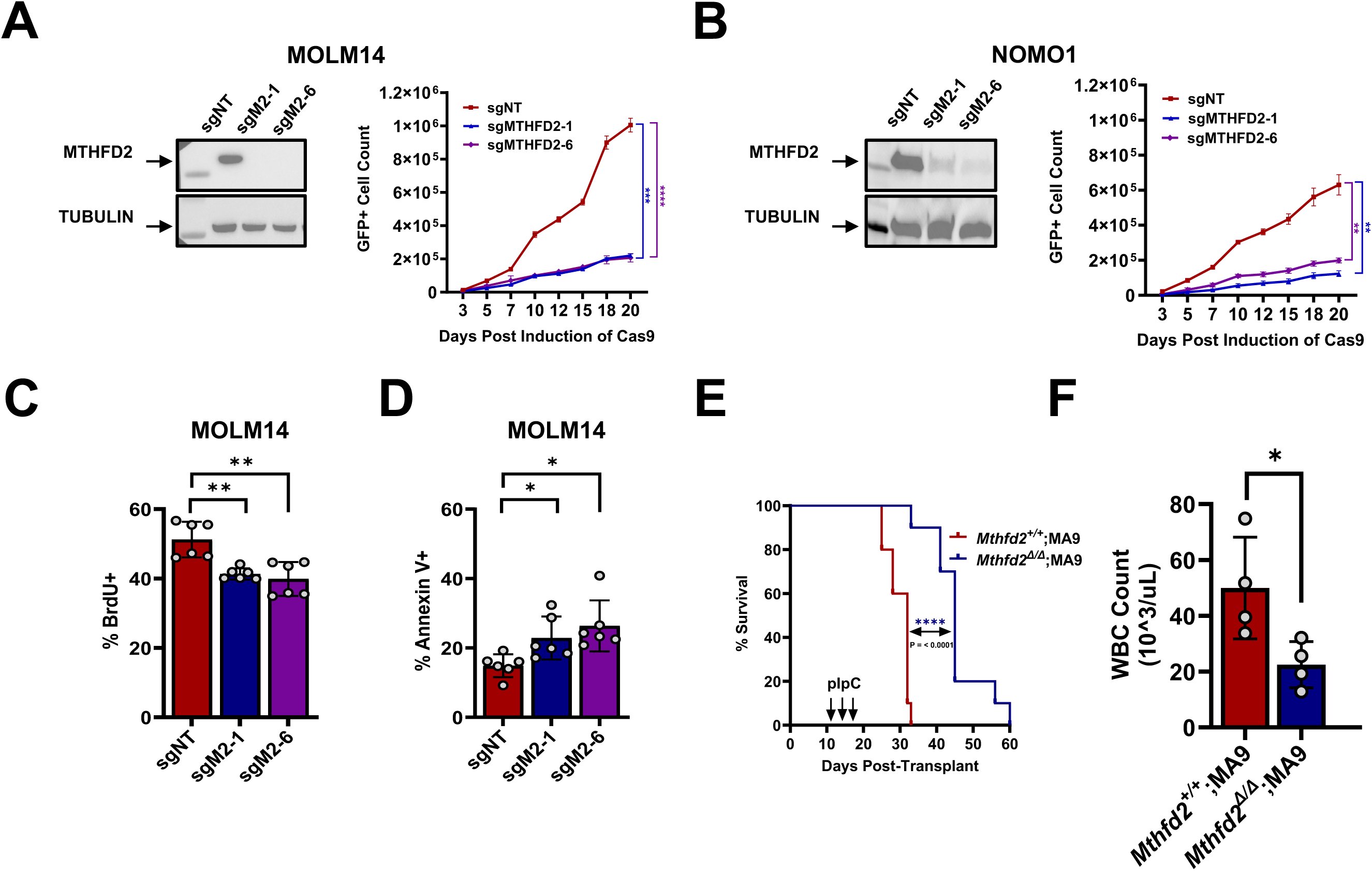
MTHFD2 Is Required for Leukemic Cell Survival and Disease Progression. **A & B.** DOX-treated MOLM14-TetON-Cas9 cells (A) and NOMO1-TetON-Cas9 cells (B) expressing sgNT, sgMTHFD2-1 or sgMTHFD2-6 were subjected to: western blot with the indicated antibodies at day four post-DOX (*left panels*) or flow cytometry to quantify total number of GFP+ cells at the indicated days post-DOX treatment (*right panels*). An unpaired t-test with Welch’s correction was used to compare the number of GFP+ cells between sgNT and each sgMTHFD2 condition (** p<0.01, ***p < 0.001, ****p < 0.0001). **C.** DOX-treated MOLM14-TetON-Cas9 cells expressing sgNT, sgMTHFD2-1 or sgMTHFD2-6 were analyzed for BrdU-incorporation at 7-days post-DOX. An unpaired t-test with Welch’s correction was used to compare the % of cells in S-phase (i.e. BrdU+) cells between sgNT and each sgMTHFD2 condition (**p < 0.01). **D**. DOX-treated MOLM14-TetON-Cas9 cells expressing sgNT, sgMTHFD2-1 or sgMTHFD2-6 were analyzed for Annexin V staining by flow cytometry at 7 days post-DOX. An unpaired t-test with Welch’s correction was used to compare the % Annexin V+ of cells between sgNT and each sgMTHFD2 condition (**p < 0.01). **E.** Mice transplanted Mthfd2^+/+^;Mx1-Cre;MA9 or Mthfd2^fl/fl^;Mx1-Cre;MA9 fully transformed leukemia BM cells were administered 3 doses of pIpC beginning at Day 10 post-transplant to induce gene deletion. Mice from each condition were then monitored for signs of AML. Kaplan-Meier analysis was applied to compare survival probabilities between Mthfd2^+/+^;MA9 or Mthfd2^Δ/Δ^;MA9 conditions and a Log-rank (Mantel-cox) test (n = 5 per group) was used to determine significance. **F.** White blood cell counts (WBC) of peripheral blood taken from Mthfd2^+/+^;MA9 and Mthfd2^Δ/Δ^;MA9 at the time of confirmed leukemia. An unpaired t-test with Welch’s correction was used to compare WBC count between Mthfd2^+/+^;MA9 and Mthfd2^Δ/Δ^;MA9 conditions (*p < 0.05).

### *Mthfd2* is dispensable for maintaining healthy hematopoiesis

We next evaluated the impact of *Mthfd2* deletion on healthy hematopoiesis (PB) (Figure 2A). Following pIpC treatment, peripheral blood (PB) from *Mthfd2*^fl/fl^;Mx1-Cre (*Mthfd2^+/+^*) and *Mthfd2^flox/flox^*;Mx1-Cre (*Mthfd2*^Δ/Δ^) mice was sampled every 4 weeks over a 20-week period and subjected to complete blood counts (CBCs) and flow cytometric analyses. Despite efficient *Mthfd2* deletion at 20 weeks post-pIpC (Supplementary Figure 2A), we did not observe any significant changes in total white blood cells (WBC), red blood cells (RBC), platelets, or lymphocytes between *Mthfd2^fl/fl^* and *Mthfd2*^Δ/Δ^ mice (Figure 2B). We also did not observe significant differences in immunophenotypic granulocytes, monocytes, T cells, or B cells between conditions (Figure 2C and Supplementary Figure 2B-2E). We also did not observe a significant difference in the number of immunophenotypic long-term hematopoietic stem cells (LT-HSCs, Lineage^low^, cKit+, Sca-1+, CD150+, CD48-, CD34-), common lymphoid progenitors (CLPs, Lineage^low^, cKit ^int^, Sca-1^int^, IL-7R+), common myeloid progenitors (CMPs, Lineage^low^, cKit+, Sca-1-, CD34^int^, FcgRII/III^int^), granulocyte-macrophage progenitors (GMPs, Lineage^low^, cKit+, Sca-1-, CD34+, FcgRII/III+), or Megakaryocyte-Erythrocyte Progenitors (MEPs, Lineage^low^, cKit+, Sca-1-, CD34-, FcgRII/III-) between the BM of *Mthfd2*^fl/fl^ and *Mthfd2*^Δ/Δ^ mice (Figure 2D).

**Figure 2.**
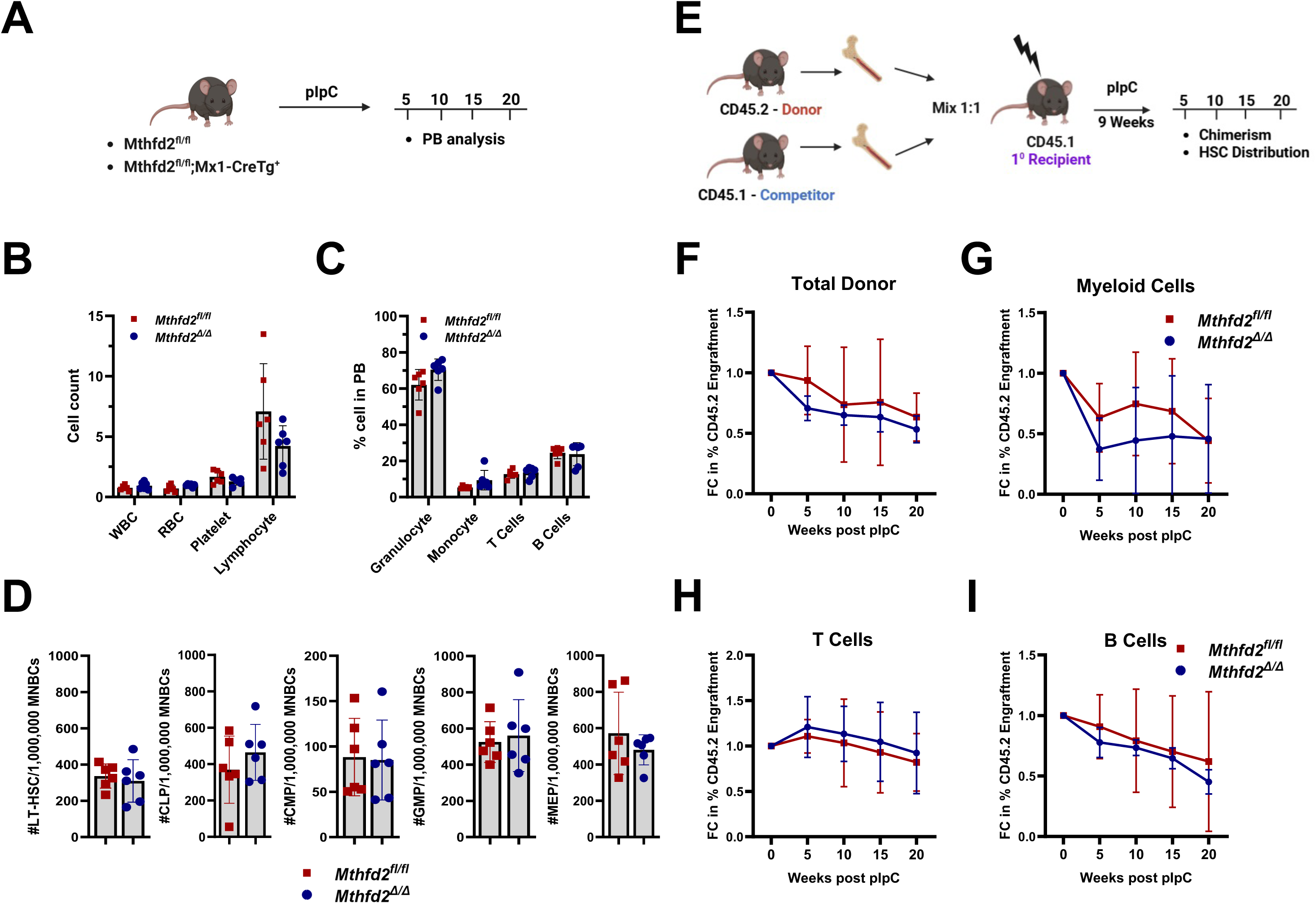
*Mthfd2* Deletion Does Not Impair Normal Hematopoiesis. **A.** Schematic of non-competitive transplantation workflow to assess mature blood cell reconstitution. **B.** Counts (10^3^/μl) of indicated PB populations between *Mthfd2^fl/fl^* and *Mthfd2^Δ/Δ^* conditions at 20 weeks post-pIpC. **C.** Flow cytometric analysis of the various mature PB hematopoietic cells between *Mthfd2^fl/fl^*and *Mthfd2^Δ/Δ^* conditions at 20 weeks post-pIpC. **D**. Flow cytometric quantification of LT-HSCs, GMPs, CMPs, and MEPs in the BM of *Mthfd2^fl/fl^* and *Mthfd2^Δ/Δ^* mice at 24 weeks post-transplant. **E.** Schematic representation of competitive primary transplantation assays to assess hematopoietic stem cell function. **F–I.** Flow cytometry analysis of PB donor-derived (CD45.2^+^) chimerism in female *Mthfd2^fl/fl^*and *Mthfd2^Δ/Δ^* primary recipients at the indicated time points post pIpC: (F) Total donor, (G) Myeloid cells, (H) T cells, (I) B cells. An unpaired t-test with Welch’s correction was used for statistical comparison of each parameter between Mthfd2^fl/fl^ and Mthfd2^Δ/Δ^ conditions.

To determine the impact of *Mthfd2* deletion on HSPC function, we performed competitive transplantation assays. Briefly, MNBCs from *Mthfd2*^fl/fl^ and *Mthfd2^fl/fl^*;Mx1Cre^+/+^ mice (both CD45.2) were separately transplanted, in competition with equal CD45.1/2 MNBCs competitor, into lethally irradiated syngeneic CD45.1 recipient mice (Figure 2E). To control for potential sex differences, parallel transplants were conducted with male donor BM being transplanted into male recipients and female donor into female recipients. Following 8-weeks of engraftment, all mice were treated with pIpC, followed by PB chimerism analysis every 4 weeks for 20 weeks. PB CD45.2 chimerism was not significantly impacted by *Mthfd2* deletion in either sex (Figure 2F and Supplementary Figure 2F). Myeloid, B cell, or T cell engraftment was also not significantly different between conditions, regardless of sex (Figure 2G-2I and Supplementary Figure 2G-2I). Collectively, these results demonstrate that loss of *Mthfd2* does not significantly impact steady-state hematopoiesis or hematopoietic reconstitution potential.

### MTHFD2 supports nucleotide synthesis in AML

Many cancer cells rely on the mitochondrial arm of one-carbon metabolism to support *de novo* synthesis of dTTP and purine^12,27^. To discern the contribution of the cytosolic and mitochondrial arms in AML, we employed a previously established isotopologue tracing assay that utilizes a deuterium-labeled serine isotopologue (2,3,3-^2^H-Serine) to quantify the contribution of each arm to dTTP synthesis (Figure 3A)^9^. Briefly, control and *MTHFD2*-deleted human AML cells were cultured in 2,3,3-^2^H-Serine-supplemented media and then extracted total polar metabolites were analyzed by hydrophilic liquid chromatography and mass spectrometry (HILIC-MS). To ensure that any observed metabolic changes were attributed to reduced *MTHFD2* expression and not to the resultant changes in cell cycle or viability, metabolites were extracted prior to the onset of cellular changes (96 hours post-transduction). Similar to many other MTHFD2-expressing cancer cell lines, AML cells primarily rely on the mitochondrial arm (M+1) to supply serine-derived carbons to dTMP and dTTP synthesis (Figure 3B and 3C). However, upon deletion of *MTHFD2*, dTMP and dTTP synthesis is primarily driven by the cytosolic arm of one-carbon metabolism (M+2) (Figure 3B and 3C). MTHFD2 inhibition had no impact on the contribution of serine to glycine synthesis (Figure 3D) but did lead to a significant decline in the utilization of serine in the *de novo* synthesis of inosine monophosphate (IMP), guanosine monophosphate (GMP) and adenosine monophosphate (AMP) (Figure 3E-3G).

**Figure 3.**
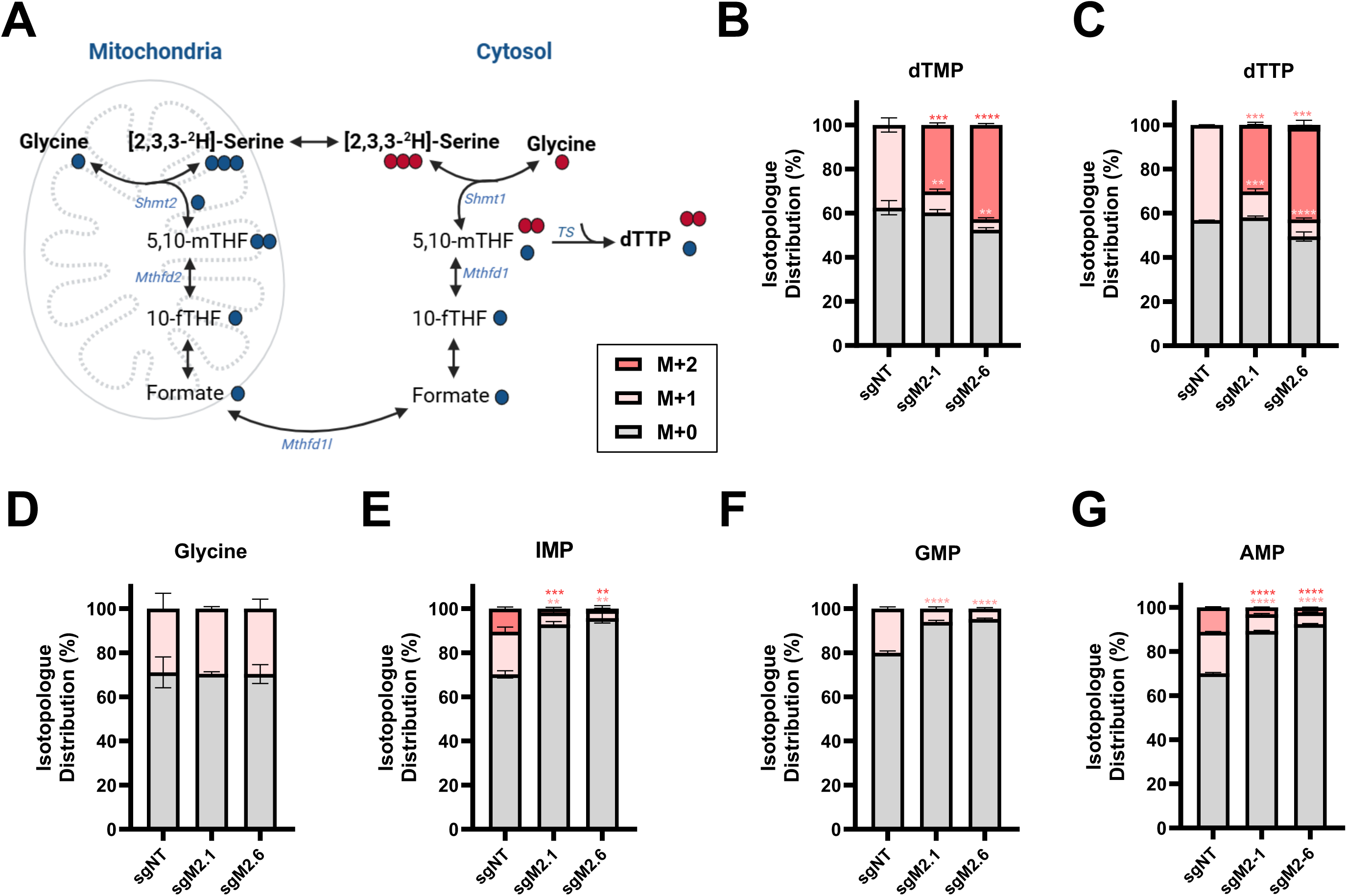
Mitochondria derived One-Carbon Metabolism is critical in AML. **A.** Schematic of [2,3,3-2H]-Serine labeling of various mitochondrial and cytosolic one-carbon metabolism intermediates. **B–G.** [2,3,3-2H]-Serine isotopologue distribution of the following one-carbon metabolites in control (sgNT) and *MTHFD2*-deleted (sgMTHFD2-1 or sgMTHFD2-6) MOLM14 cells: (B) dTMP, (C) dTTP, (D) Glycine, (E) IMP, (F) GMP, (G) AMP. Unpaired, t-test with Welch’s correction were used to compare M+1 or M+2 labeling of each metabolite between sgNT and each sgMTHFD2 condition (** p<0.01, ***p < 0.001, ****p < 0.0001).

As an orthogonal approach to assess the impact of *MTHFD2* loss on one-carbon metabolism, we performed metabolic tracing using a uniformly ^13^C-labeled serine isotopologue (U-^13^C-Serine) (Supplementary Figure 3A). MTHFD2 inhibition did not impact serine-derived carbons being shunted into glycine or dTTP synthesis (Supplementary Figure 3B and 3C), confirming that the cytosolic arm of one-carbon metabolism can maintain dTTP synthesis in the absence of MTHFD2. Loss of *MTHFD2* expression also significantly impeded *de novo* synthesis of AMP and GMP using this carbon-based isotopologue (Supplementary Figure 3D and 3E).

To determine whether the decline in *de novo* purine synthesis was contributing to the anti-leukemia effects of *MTHFD2* deletion, we cultured control and *MTHFD2*-deleted human AML cells in the presence or absence of exogenous purines (deoxyadenosine and deoxyguanosine; dP). Purine supplementation partially restored the expansion of *MTHFD2*-deleted leukemia cells (Supplementary Figure 3F and 3G), suggesting that purine synthesis, in part, contributed to the anti-leukemia effects of *MTHFD2* loss.

### MTHFD2 is required to maintain mitochondrial NADH and NADPH levels

MTHFD2 is a bifunctional enzyme that converts 5,10-dimethylene-tetrahydrofolate (5,10-me-THF) into 10-formyl-tetrahydrofolate (10-fTHF) via a two-step reaction. MTHFD2 first dehydrogenates 5,10-me-THF into 5,10-methylenyl-tetrahydrofolate and then converts this species into 10-fTHF^13^. The hydrogen atom removed from 5,10-me-THF is used to reduce NAD^+^ to NADH or NADP^+^ to NADPH. The MTHFD2-mediated reduction of NAD^+^ or NADP^+^ can be indirectly determined via 2,3,3-^2^H-Serine tracing^9^. Specifically, the conversion of oxaloacetic acid (OAA) into malic acid utilizes MTHFD2-generated NADH; therefore, levels of serine-derived hydrogens in malate can be used to indirectly quantify changes in MTHFD2-generated NADH^9^ (Figure 4A). Additionally, NADPH generated by MTHFD2 is used to convert γ-glutamyl phosphate into proline, allowing for proline serine-derived hydrogens to be used as an indirect measure of MTHFD2-mediated NADPH production^9^ (Figure 4B). Compared to controls, *MTHFD2*-deleted cells display significantly less serine-derived deuterium incorporation into both malate and proline (Figure 4A and 4B), indicating that MTHFD2 disrupts NADH and NADPH production in the mitochondria.

**Figure 4.**
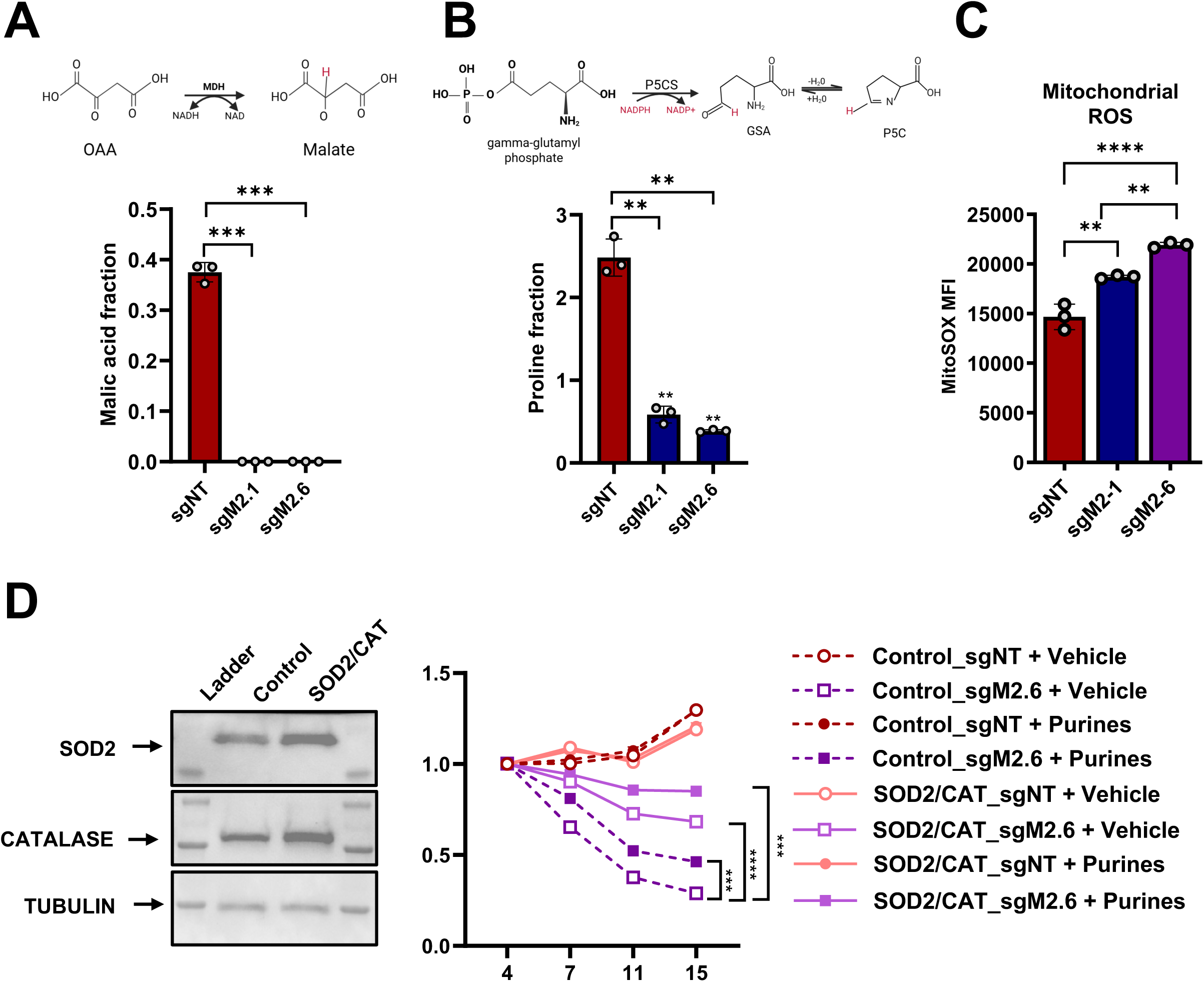
MTHFD2 Is Crucial for Maintaining Redox Homeostasis In AML. **A–B.** [2,3,3-2H]-Serine isotopologue distribution of the following one-carbon metabolites in control (sgNT) and *MTHFD2*-deleted (sgMTHFD2-1 or sgMTHFD2-6) MOLM14 cells: (A) Malic acid and (B) Proline. The top panels represent a schematic of each respective reaction, and the bottom panels are the graphical representation of the detected isotopologue labeled malic acid (B) and proline. Unpaired t-test with Welch’s correction were used to compare M+1 or M+2 labeling of each metabolite between sgNT and each sgMTHFD2 condition (** p<0.01, ***p < 0.001, ****p < 0.0001). **C.** Control (sgNT) and *MTHFD2*-deleted MOLM14 cells were analyzed for MitoSOX staining by flow cytometry at four days post-MTHFD2 deletion. An unpaired, nonparametric t*-*test was used to compare the MFI of MitoSOX between sgNT and each sgMTHFD2 conditions or between the two sgMTHFD2 conditions (**p < 0.01, ***p < 0.0001). **D.** Control (sgNT) and *MTHFD2*-deleted (sgMTHFD2-1 or sgMTHFD2-6) MOLM14 cells were engineered to co-express BFP either alone (Control) or superoxide dismutase and catalase, in tandem (SOD2/CAT). Cells from each condition were treated with vehicle or 50 μM of deoxypurines and then monitored for fold change in BFP over time (compared to Day 4). An unpaired t-test with Welch’s correction was used to compare each indicated two-condition comparison (***p < 0.001, ****p < 0.0001).

### AML cells rely on MTHFD2 to maintain mitochondrial redox homeostasis

NADPH plays a key role in maintaining mitochondrial redox homeostasis^28^. Therefore, we hypothesized that disruption of MTHFD2-mediated NADPH generation may lead to mitochondrial ROS imbalances. In fact, *MTHFD2*-deletion resulted in significantly higher steady-state levels of mitochondrial ROS compared to controls (Figure 4C). To assess whether the induction of mitochondrial ROS was contributing to the anti-leukemia effects of MTHFD2 inhibition, we engineered AML cells to overexpress SOD2 and Catalase (CAT), two enzymes that neutralize mitochondrial superoxide when expressed in tandem^29^. Overexpression of SOD2 and CAT partially protected AML cells from *MTHFD2* deletion, confirming that MTHFD2, in part, regulates mitochondrial ROS homeostasis to support AML cell biology (Figure 4D and Supplementary Figure 4A-4C). We next examined how the combination of purine supplementation and ROS-neutralization impacted AML cells. Remarkably, the combination of SOD2-CAT overexpression and exogenous purines supplementation was superior to each intervention alone and almost completely reversed the anti-leukemia effects of MTHFD2 inhibition (Figure 4D and Supplementary Figure 4B and 4C). These results suggest that AML cells rely on MTHFD2 to support *de novo* purine synthesis and maintain mitochondrial redox homeostasis.

### Pharmacological inhibition of mitochondrial MTHFD2 displays anti-leukemia activity

Bonagas *et al.,* previously reported the development of a chemical inhibitor of MTHFD2, TH9619, that displayed anti-leukemia effects in experimental models of AML^22^. In a follow-up study, it was shown that TH9619 is unable to inhibit mitochondrial MTHFD2 but rather inhibits nuclear MTHFD2 as well as cytosolic MTHFD1^23^. Since our results indicated that mitochondrial MTHFD2 supports AML cell biology, we sought to identify a chemical inhibitor capable of impeding mitochondrial MTHFD2 activity. We focused on DS18561882, an MTHFD2 inhibitor that demonstrates *in vivo* anti-tumor efficacy in a breast cancer xenograft model^30^. To assess the capacity of DS18561882 to block mitochondrial MTHFD2 activity, we treated AML cells with either vehicle or DS18561882, and then performed 2,3,3-^2^H-Serine tracing. DS18561882 phenocopied *MTHFD2* deletion by effectively blocking mitochondrial serine from contributing to dTTP and purine synthesis as well as MTHFD2-mediated NADPH generation (Figure 5A-5E). Conversely, TH9619 did not significantly impact the mitochondrial arm of one-carbon metabolism (Figure 5F-5J).

**Figure 5.**
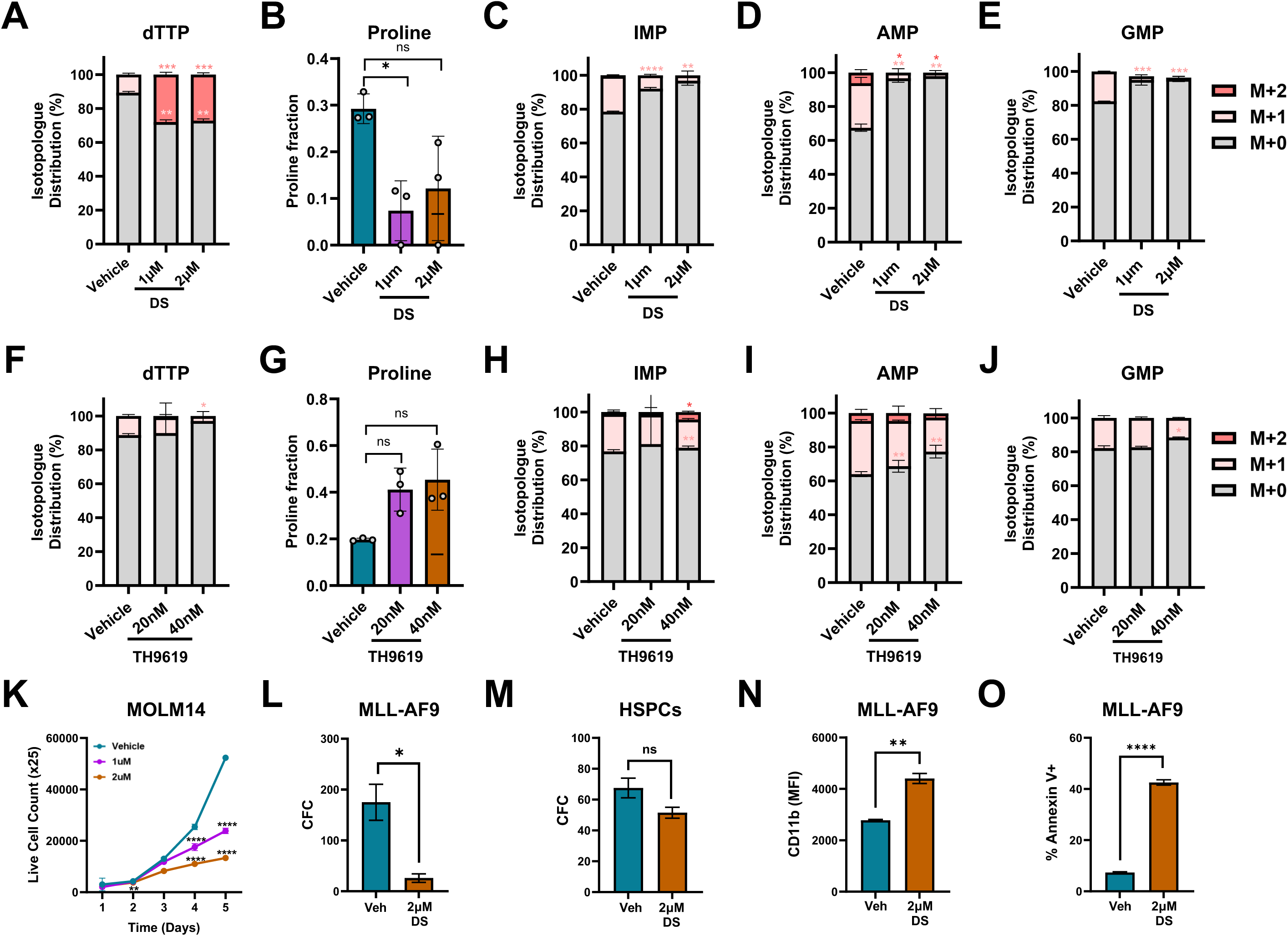
DS18561882, but Not TH9619, Blocks Mitochondrial One-Carbon Metabolism in AML Cells. **A–E.** [2,3,3-2H]-Serine isotopologue distribution of the following one-carbon metabolites in Vehicle and DS18561882-treated (1µM and 2µM) treated MOLM14 cells: (A) dTTP, (B) Proline, (C) IMP, (D) AMP, (E) GMP. Unpaired t-test with Welch’s correction were used to compare M+1 or M+2 labeling of each metabolite between Vehicle and DS18561882-treated MOLM14 cells conditions (* p<0.01, ** p<0.01, ***p < 0.001, ****p < 0.0001). **F–J.** [2,3,3-2H]-Serine isotopologue distribution of the following one-carbon metabolites in Vehicle and TH9619-treated (20nM and 40nM) MOLM14 cells: (F) dTTP, (G) Proline, (H) IMP, (I) AMP, (J) GMP. Unpaired t-test with Welch’s correction were used to compare M+1 or M+2 labeling of each metabolite between Vehicle and TH9619-treated MOLM14 cells conditions (* p<0.01, ** p<0.01). **K**. Growth curve showing change in viability following treatment of MOLM14 AML cells with DS18561882. **L-M.** Change in colony count following treatment with DS18561882 in **L.** MLL-AF9 cells **M.** healthy HSPCs **N.** MLL-AF9 cells were treated with either vehicle or 2µM DS18561882 were analyzed for CD11b MFI by flow cytometry at 96 hours post-treatment. An unpaired t-test with Welch’s correction was used to compare the CD11b MFI of cells between vehicle or 2µM DS18561882 condition (**p < 0.01). **O.** MLL-AF9 cells were treated with either vehicle or 2µM DS18561882 were analyzed for Annexin V staining by flow cytometry at 4 days post-treatment. An unpaired t-test with Welch’s correction was used to compare the percentage of Annexin V positive cells between vehicle or 2µM DS18561882 condition (***p < 0.0001).

We next tested the impact of DS18561882 treatment across a panel of mouse and human AML cell lines. DS18561882 reduced the viability of the human AML cell lines MOLM-14, OCI-AML3 and NOMO-1, in a dose-dependent manner (Figure 5K and Supplementary Figure 5A and 5B). THP-1 and K562 cells were also significantly impeded by DS18561882 treatment (Supplementary Figure 5C and 5D). Similarly, increasing concentrations of DS18561882 reduced the viability of mouse AML cells expressing either MLL-AF9 or a combination of Dnmt3a^R878H^ and FLT3-ITD (DF cells) (Supplementary Figure 5E and 5F). Consistent with genetic deletion of MTHFD2, DS18561882 treatment significantly reduced the colony-forming capacity (CFC) of MLL-AF9-expressing AML cells, but not that of healthy HSPCs (Figure 5L and 5M). DS18561882-treated mouse MLL-AF9-expressing AML cells also displayed increased CD11b expression and cell death compared to controls (Figure 5N and 5O). Reinforcing that DS18561882 can bind to and inhibit mitochondrial MTHFD2, DS18561882 treatment increased levels of mitochondrial ROS (Supplementary Figure 5G). Furthermore, the mitochondria-targeted antioxidant that scavenges superoxide^29^ suppressed the cytotoxic effects of DS18561882 (Supplementary Figure 5H).

Given that AML comprises numerous genetically distinct subtypes, we next assessed the anti-leukemia activity of DS18561882 across a genetically diverse panel of PD-AML samples. Sixty bone marrow-derived PD-AML samples were treated with increasing concentrations of DS18561882 and then assessed for viability. DS18561882 treatment led to a dose-dependent decline in viable cell numbers across PD-AML samples with varying responses (Figure 6A). In contrast, bone marrow–derived hematopoietic cells from three healthy donors showed minimal changes in viability upon treatment with DS18561882 (Figure 6B).

**Figure 6.**
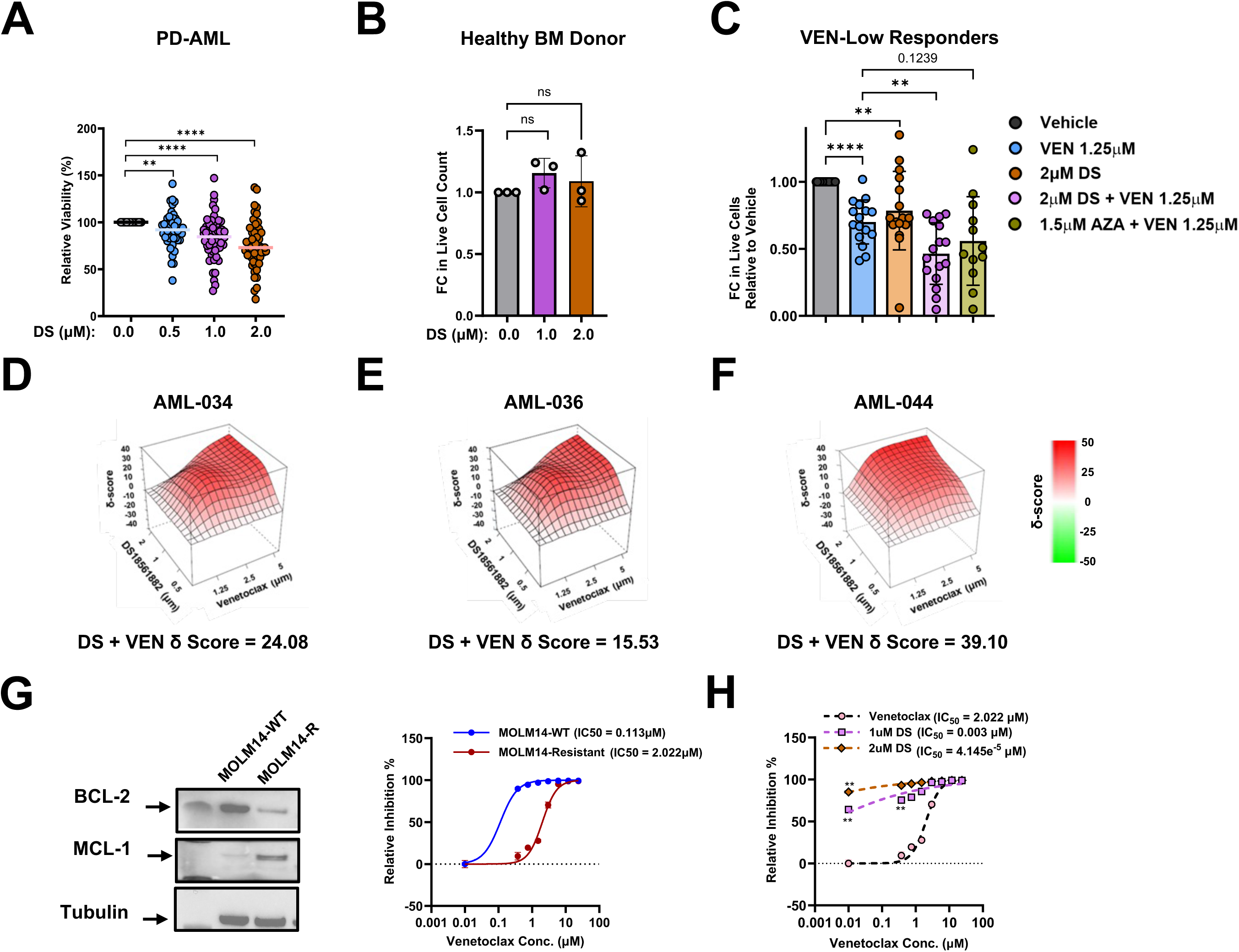
Pharmacologic inhibition of MTHFD2 selectively impairs AML viability while sparing normal hematopoietic cells. **A.** Change in cell viability relative to vehicle in patient-derived AML cells following treatment with 0.5, 1 or 2µM DS18561882 for 96 hours. An unpaired, nonparametric *t-*tests were used to compare control and treated groups (** p<0.01, ****p < 0.0001). **B.** Change in cell viability in healthy patient derived bone marrow cells following treatment with DS18561882 for 96 hours. **C.** Comparison of response to either 2 µM DS18561882 alone, VEN alone or VEN and AZA in Venetoclax resistant (VEN-Low Responders) PD-AML cell lines. Unpaired, nonparametric *t-*tests were used to compare control and treated groups (** p<0.01, ****p < 0.0001). **D-F.** Three-dimensional plots of the synergy δ scores for select PD-AML samples treated with varying combinations of the DS18561882 (x-coordinate), and VEN (y-coordinate). **G.** MOLM14 parental cells (WT) and resistant cells (R) were subjected to western blot with the indicated antibodies (*left panels*). Dose response curves comparing IC₅₀ in MOLM14 WT and Resistant cells at 72hours of treatment with either Venetoclax or vehicle *(right panel).* **H.** Dose response curves comparing IC₅₀ in MOLM14-R cells following treatment with either vehicle, 1 µM or 2 µM DS18561882 (**p < 0.01).

We next compared the activity of DS18561882 with genetic subtype across PD-AML samples for which paired genetic data was available. Samples were stratified into two catergories: 1) DS-High Responders, defined as those exhibiting cell viability below the cohort median following treatment with 2 µM DS18561882; and 2) DS-Low Responders, defined as those with viability above the median (Supplementary Figure 6A). Notably, 5 of 6 PD-AML samples bearing *IDH2* mutations were characterized as DS-High Responders (i.e. those PD-AML cells that were more sensitive to DS18561882) (Supplementary Figure 6A and 6B). Additionally, 4 of 5 samples with a *KIT*, *RUNX1* or *ASXL1* mutation were enriched in DS-High Responders (Supplementary Figure 6C-6E). In contrast, all PD-AML samples harboring *KRAS* mutations (n = 3) were classified as DS-Low Responders (Supplementary Figure 6F). Although not statistically significant, the majority of samples bearing a *NPM1* or *NRAS* mutant (60% and 56%, respectively) demonstrated lower responses to DS18561882 (Supplementary Figure 6A).

### MTHFD2 inhibition and Venetoclax display cooperative anti-leukemia activity

We next investigated whether DS18561882 treatment cooperated with the FDA-approved agents Cytarabine (Ara-C), Doxorubicin (DOXO), 5-azacytidine (AZA), or Venetoclax. We did not observe any cooperation between DS18561882 and either Ara-C or DOXO (Supplementary Figure 7A-7C). However, DS18561882 cooperated with both AZA and Venetoclax treatment, though the relationship between DS18561882 and Venetoclax was the stronger of the two combinations (Supplementary Figure 7A, 7D, and 7E). Genetic deletion of MTHFD2 also sensitized human AML cells to Venetoclax treatment (Supplementary Figure 7F), and the combination of Venetoclax and MTHFD2 inhibition, genetically or pharmacologically, increased mitochondrial ROS generation compared to each condition alone (Supplementary Figure 8A-8D).

Venetoclax is a BCL-2 inhibitor that has emerged as a promising therapy for certain subgroups of AML patients, though resistance remains a clinical obstacle^31,32^. Therefore, we assessed how the combination of DS18561882 and Venetoclax impacted PD-AML viability. For this analysis, 31 PD-AML samples were treated with increasing concentrations of either DS18561882 or Venetoclax alone as well as in combination. Because Venetoclax is used in combination with AZA clinically^31^, we included this treatment as a benchmark in 23 PD-AML samples. Similar to patients, we observed that PD-AML samples differentially responded to Venetoclax treatment alone. Specifically, we found that 15 of the 31 PD-AML samples displayed a 70% or greater reduction in cell viability between Venetoclax and vehicle treatment conditions, and we therefore categorized these samples as VEN-High Responders (VEN-High-Resp.) and the other 16 PD-AML samples as VEN-Low Responders (VEN-Low-Resp.) (Supplementary Figure 9A). The 16 VEN-Low Responder samples were enriched for mutations previously linked to Venetoclax resistance (*FLT3, TP53, ASXL1*, and spliceosome genes (*SRSF2,* and *U2AF1*) (Supplementary Figure 9B)^33,34^. Both VEN-High-Responder and VEN-Low-Responder PD-AML samples displayed a marginal but significant decrease in viability upon treatment with 1 µM or 2 µM DS18561882. Although Venetoclax alone was highly effective in reducing the mean viability of VEN-High-responder samples (14.4%), the addition of 1 µM or 2 µM DS18561882 further reduced viability to 10% and 9%, respectively (Supplementary Figure 9C and 9D). Moreover, the combination of venetoclax and either 1 µM or 2 µM DS18561882 was significantly more effective than Venetoclax alone in reducing the viability VEN-Low Responder samples (Figure 6C and Supplementary Figure 9E). While the addition of AZA improved the cell killing effects of Venetoclax, this combination was not significantly better than Venetoclax treatment alone in reducing the viability of VEN-Low Responder samples (Figure 6C and Supplementary Figure 9E).

Using SynergyFinder^35^, we calculated the synergy scores (δ score) for 19 of the 31 PD-AML samples treated with combination therapies. We observed that 9/19 PD-AML (47%) displayed synergistic cooperation between Venetoclax and DS18561882 whereas 5/19 PD-AML (26%) displayed a synergistic cooperation between Venetoclax and AZA (Figure 6D-F and Supplementary Table 1). Notably, the 9 samples that displayed synergy between Venetoclax and DS18561882 were all samples that responded poorly to Venetoclax treatment alone (Supplementary Table 1). We did not observe any significant relationship between the combined effectiveness of Venetoclax and DS18561882 with a specific AML mutation.

To evaluate the impact of DS18561882 treatment on AML cells that had developed Venetoclax resistance, we generated Venetoclax-resistant cells (MOLM14-R) through chronic exposure to increasing concentrations of Venetoclax as shown previously^36^. Relative to parental controls (MOLM14-WT), MOLM14-R cells exhibited over 17-fold resistance to Venetoclax and displayed elevated expression of MCL-1, a common mechanism of Venetoclax resistance in AML^32,36^ (Figure 6G). Venetoclax-resistant cells displayed heightened sensitivity to 1 and 2 µM DS18561882 monotherapy, shifting the IC₅₀ from 2.022 µM in control conditions to 0.003 µM and 4.15e^−5^, respectively (Figure 6H). Similar to DS18561882 treatment, MOLM14-R cells were more sensitive to *MTHFD2* deletion compared to the vehicle control (Supplementary Figure 9F).

Collectively, these data suggest that MTHFD2 drives purine synthesis and maintains mitochondrial redox homeostasis to support AML, and that pharmacological targeting of MTHFD2 in combination with existing AML therapies, such as Venetoclax, may have therapeutic value.

## Discussion

Previous studies showing that genetic inhibition of MTHFD2 or small molecules that co-target nuclear MTHFD2 and cytosolic MTHFD1 antagonize the AML growth, have positioned MTHFD2 as a potential therapeutic target in AML^11,21–23^. Further strengthening this position, we report here that deletion of MTHFD2 inhibits AML cell growth and survival both *in vitro* and *in vivo* without impacting healthy HSPC function. AML cells have been shown to rely on MTHFD2 in the nucleus to regulate replication fork progression. However, these results are based on an inhibitor that effectively blocks nuclear and cytoplasmic MTHFD2 but is unable to penetrate the mitochondria. Thus, the mechanism by which MTHFD2 supports AML remains incomplete. Using a combination of genetic ablation and an MTHFD2 inhibitor that is able to block mitochondrial MTHFD2 activity, we report that, in addition to replication fork activity, AML cells rely on MTHFD2 to support *de novo* purine synthesis and maintain mitochondrial redox status. We also show that neither genetic ablation nor pharmacological inhibition of MTHFD2 significantly impacted healthy hematopoietic cells. Previous reports have shown that leukemic cells exhibit higher levels of replication and mitochondrial ROS compared to HSPCs^37,38^, possibly explaining why AML cells are more reliant on MTHFD2 than healthy hematopoietic cells.

AML is a genetically heterogeneous disease with distinct subtypes that influence prognosis and guide treatment decisions.^18,20,39^. Although the precise genetic subtypes that are particularly reliant on MTHFD2 remain to be determined, our treatment of a cohort of PD-AML samples revealed that certain AML recurring mutations associate with differential responses to MTHFD2 inhibition. For example, we found that PD-AML samples bearing *IDH2*, *ASXL1*, *KIT*, or *RUNX1* mutations tended to be more responsive to DS18561882. Notably, each of these mutations have been previously associated with elevated ROS production. Specifically, IDH mutations show a heightened susceptibility to mitochondrial-targeting agents in cancer cells^40–42^. Furthermore, IDH2 mutations produce the oncometabolite D-2-hydroxyglutarate, which perturbs mitochondrial function and increases NAPDH consumption^43^. RUNX1 or ASXL mutation also associate with increased ROS in a PI3K/AKT-dependent manner whereas KIT mutations are associated with increased mitochondria number and ROS^44–46^. Thus, these mutations may cause AML cells to be more reliant on MTHFD2 for NADPH generation and redox buffering. Although oncogenic *KRAS* has been linked to mitochondrial one-carbon metabolism^47,48^, the three *KRAS*-mutant AML samples in our cohort were less responsive to DS18561882. Those samples also carry additional recurrent mutations; thus it is possible that these mutations may provide metabolic compensation. Alternatively, *RAS* mutations can reprogram metabolic pathways in AML and are associated with adaptive signaling that can reduce sensitivity to therapeutic agents^49^. Furthermore, oncogenic signaling, including *KRAS*, orchestrates broad metabolic rewiring – such as nucleotide and redox pathways – to influence therapeutic response and contribute to resistance mechanisms, including resistance to Venetoclax in AML^47,48,50–52^. It is important to note that given the modest size of our patient cohort and the frequent co-occurrence of mutations in AML, these findings should be interpreted as enrichment patterns that require future studies in genetically controlled experimental systems, in order to determine the relationship between MTHFD2 and certain AML genotypes.

In addition to evaluating DS18561882 as a single agent, we also explored combinations with several FDA-approved AML therapies. While DS18561882 did not cooperate with Ara-C or DOXO, we did find that the genetic or pharmacological inhibition of MTHFD2 enhanced the cytotoxic effects of Venetoclax. The addition of DS18561882 was most effective in enhancing Venetoclax activity on AML samples that are less sensitive to Venetoclax treatment alone. Previous studies have shown that the anti-leukemia activity of Venetoclax is due to, at least in part, the excessive production of mitochondrial ROS production^53–56^. We have observed that the combination of Venetoclax and MTHFD2 inhibition leads to an accumulation of mitochondrial ROS that exceeds that of each intervention alone, providing a possible explanation for their selective cooperation. We also observed that AML cells with established resistance to Venetoclax are particularly sensitive to DS18561882 treatment or MTHFD2 deletion. In the setting of Venetoclax-resistance, AML cells may utilize multiple redox buffering systems, including MTHFD2, to maintain mitochondrial redox homeostasis. However, additional studies using patient samples that have developed Venetoclax-resistance will be needed to confirm this possibility.

There are two additional points from our study that must be considered. First, although DS18561882 has a higher selectivity for MTHFD2, DS18561882 can bind and inhibit MTHFD1 and therefore, off target inhibition of MTHFD1 could be contributing to the anti-leukemia effects of DS18561882^57^. In fact, the anti-leukemia activity of another MTHFD2 inhibitor, TH91619 stems from the combined inhibition of nuclear MTHFD2 and cytosolic MTHFD1^22,23^. However, we did observe a significant phenotypic overlap between DS18561882 treatment and selective MTHFD2 deletion. Second, in addition to venetoclax, we did observe that DS18561882 and AZA showed a degree of cooperation in reducing AML cell viability. These data warrant the evaluation of combining Venetoclax, AZA and DS18561882 in the treatment of AML, however, toxicity studies on healthy tissue will be needed in parallel.

## Conclusion

We have identified MTHFD2 as a metabolic dependency that support AML by sustaining nucleotide biosynthesis and redox balance but is dispensable for healthy HSPC function. We also demonstrate that MTHFD2 inhibition cooperates with Venetoclax to restore sensitivity in Venetoclax-refractory disease. These results support preclinical rationale for further investigation of mitochondrial MTHFD2-targeting strategies for potential incorporation into AML therapeutic regimens, such as the combination of Venetoclax and Azacytidine.

**Materials and Methods** are located in Supplementary File 1

## Supporting information

3_SUPPLEMENTARY_FIGURES_JSSMS31626

4_SUPPLEMENTARY_FILE_JSSMS31626

