## Supplementary material for "Targeting MTHFD2 disrupts mitochondrial redox homeostasis and restores venetoclax sensitivity in acute myeloid leukemia": 3_SUPPLEMENTARY_FIGURES_JSSMS31626

### Supplementary Figure 1

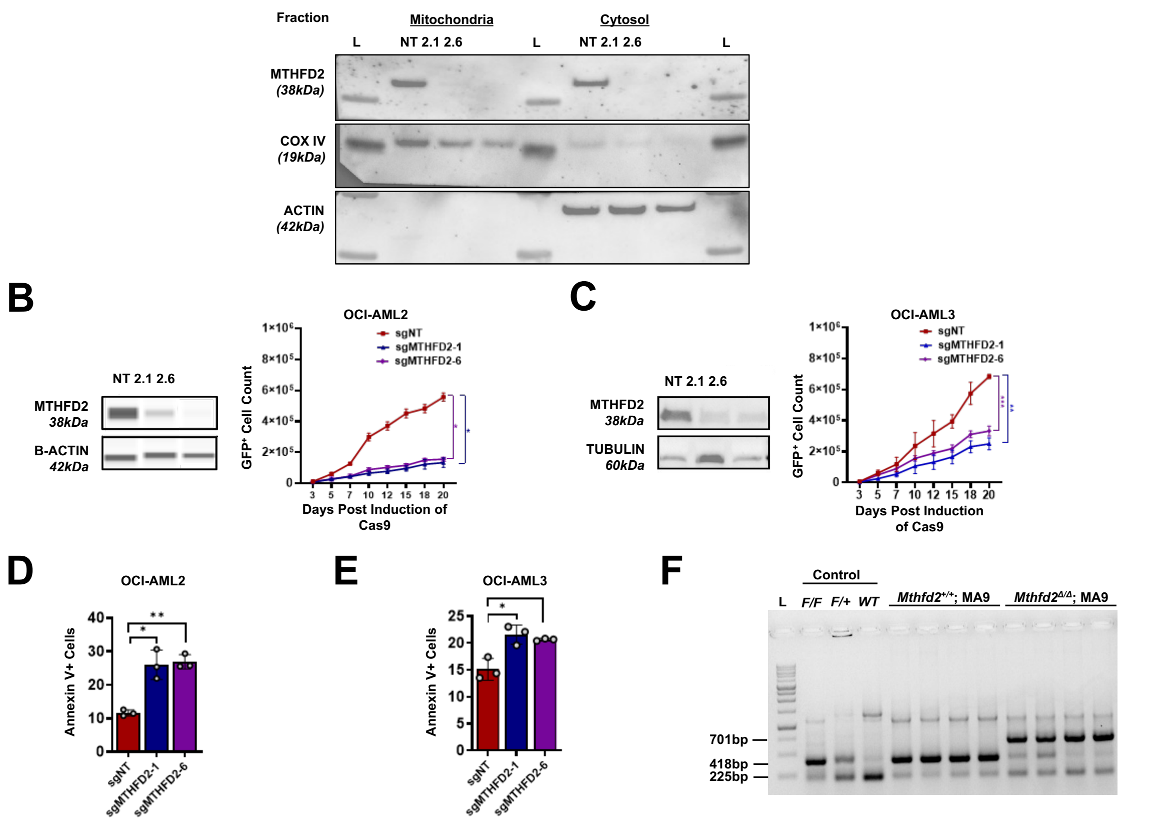


**Supplementary Figure 1. A.** Fractionation analysis of MOLM14 cells expressing sgNT, sgMTHFD2-1, or sgMTHFD2-6. **B & C.** DOX-treated OCIAML2-TetON-Cas9 cells (B) and OCIAML3-TetON-Cas9 cells (C) expressing sgNT, sgMTHFD2-1 or sgMTHFD2-6 were subjected to: western blot with the indicated antibodies at day four post-DOX (*left panels*) or flow cytometry to quantify total number of GFP+ cells at the indicated days post-DOX treatment (*right panels*). An unpaired t-test with Welch's correction was used to compare the number of GFP+ cells between sgNT and each sgMTHFD2 condition (** p<0.01, ***p < 0.001, ****p < 0.0001). **D & E.** DOX-treated OCIAML2-TetON-Cas9 cells (D) and OCIAML3 (E) expressing sgNT, sgMTHFD2-1 or sgMTHFD2-6 were analyzed for Annexin V staining by flow cytometry at 7 days post-DOX. An unpaired t-test with Welch's correction was used to compare the % Annexin V+ of cells between sgNT and each sgMTHFD2 condition (**p < 0.01, **p < 0.001). **F.**

Excision PCR analysis of BM cells from *Mthfd2*$^{+/+}$;MA9 and *Mthfd2*$^{Δ/Δ}$;MA9 secondary transplant mice.

### Supplementary Figure 2

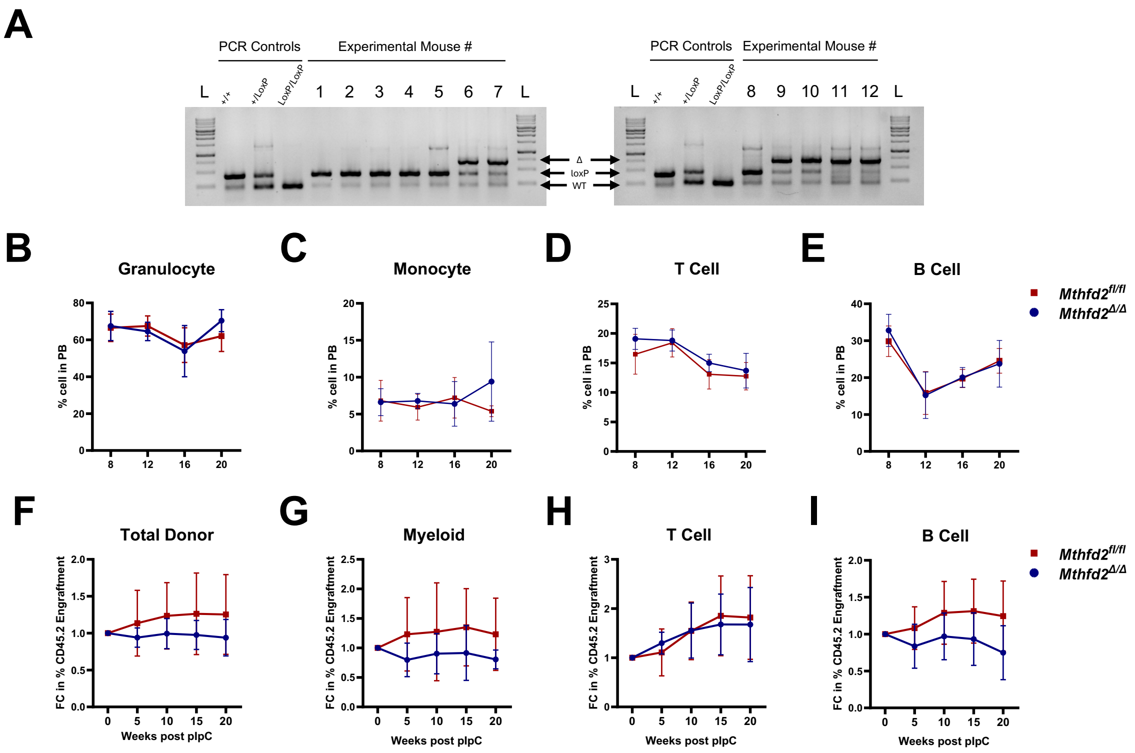


**Supplementary Figure 2. A.** Excision PCR analysis of BM cells from non-competitive transplant mice at week 24 post-pIpC. PCR amplification bands = wild type allele (WT, 225 bp), floxed allele (loxP, 418 bp), and excised allele (Δ, 701 bp). **B-E.** Flow cytometric quantification of (B) Granulocyte, (C) Monocyte, (D)**.** T cells, (E) B cells between *Mthfd2*$^{fl/fl}$ and *Mthfd2*$^{\Delta/\Delta}$ mice, **F-I.** Flow cytometry analysis of PB donor-derived (CD45.2$^{+}$) chimerism in male *Mthfd2*$^{fl/fl}$ and *Mthfd2*$^{\Delta/\Delta}$ primary recipients at the indicated time points post pIpC: (F) Total donor, (G) Myeloid cells, (H) T cells, (I) B cells. An unpaired t-test with Welch's correction was used for statistical comparison of each parameter between Mthfd2$^{fl/fl}$ and Mthfd2$^{\Delta/\Delta}$ conditions.

### Supplementary Figure 3

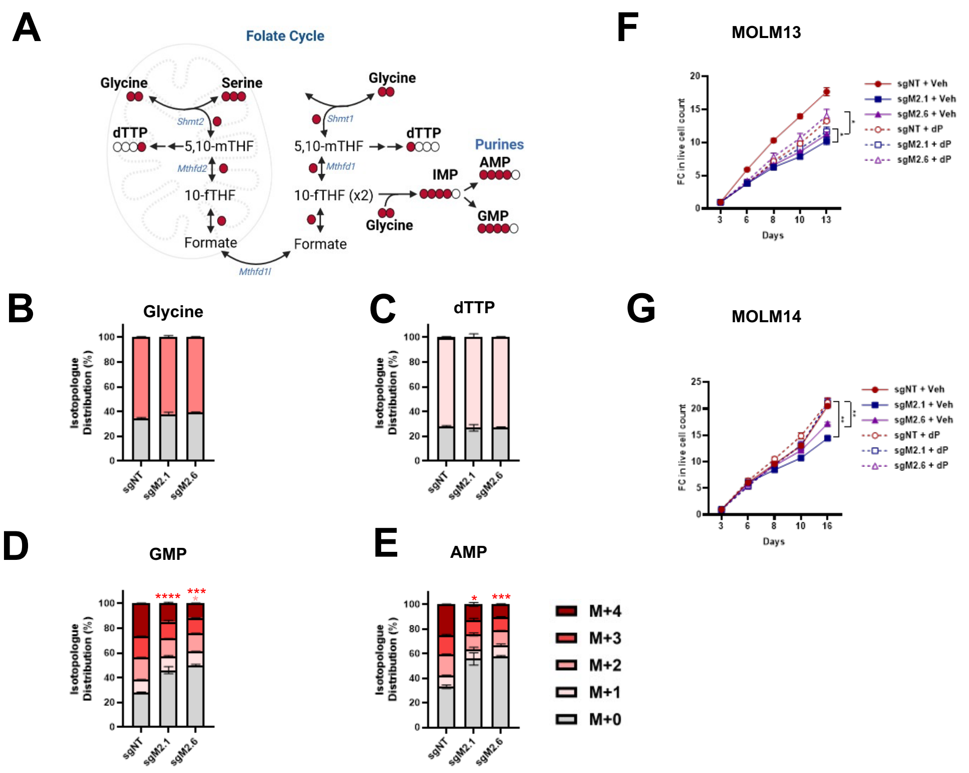


**Supplementary Figure 3**. **A.** Schematic of serine-derived carbon flux through the folate cycle. **B-E.** Serine isotopologue distribution of the following one-carbon metabolites in control (sgNT) and MTHFD2-deleted (sgMTHFD2-1 or sgMTHFD2-6) MOLM14 cells: (B) Glycine, (C) dTTP, and (D) GMP (E) AMP in sgMTHFD2 cells. An unpaired t-test with Welch's correction was used to compare isotopologue distribution between sgNT and each sgMTHFD2 condition (* $p<0.05$, ***$p < 0.001$, ****$p < 0.0001$). **F-G.** Cells from either control (sgNT) and MTHFD2-deleted (sgMTHFD2-1 or sgMTHFD2-6) were treated with vehicle or 50 μM of deoxypurines (dP) and then monitored for fold change in GFP over time (compared to Day 3) in MOLM13 (F) and MOLM14 (G) cells. An unpaired t-test with Welch's correction was used to compare each indicated two-condition comparison (*$p < 0.05$, **$p < 0.01$).

### Supplementary Figure 4

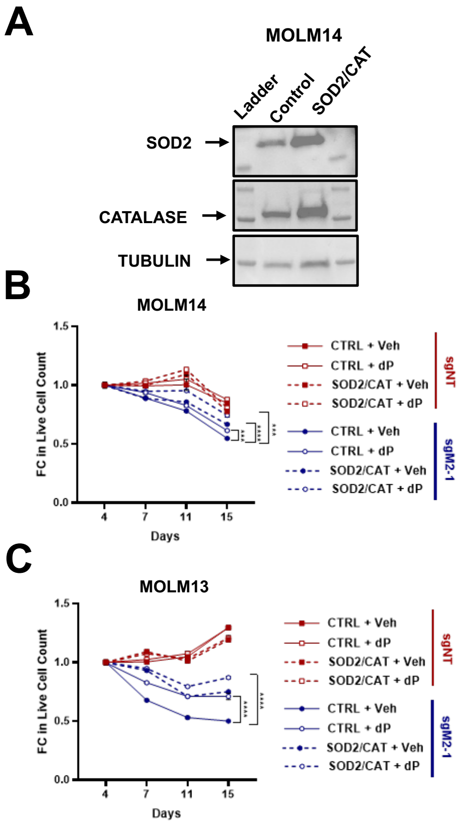


**Supplementary Figure 4. A.** MOLM14 cells expressing sgNT, sgMTHFD2-1 or sgMTHFD2-6 were subjected to western blot with the indicated antibodies **(B-C).** MOLM14 cells expressing sgNT, sgMTHFD2-1 or sgMTHFD2-6 were subjected to flow cytometry to quantify total number of live cells in control or sgMTHFD2-deleted cells expressing control or SOD2/CAT at the indicated days post-MTHFD2 deletion in MOLM14 (B) and MOLM13 (C) AML cells. An unpaired t-test with Welch's correction was used to compare the change in live cells between each indicate two-condition comparison (** p<0.01, ***p < 0.001, ****p < 0.0001).

### Supplementary Figure 5

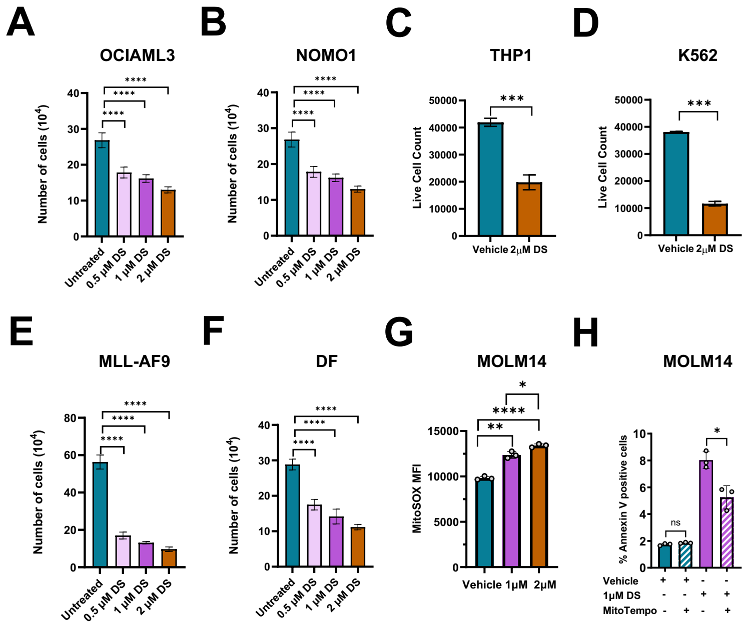


**Supplementary Figure 5. A-F.** Flow cytometry analysis to quantify total number of live cells in (A) OCIAML3 (B) NOMO1, (C) MOLM14, (D) THP1, (E) K562 AML cells following treatment with DS18561882 for 96 hours. An unpaired t-test with Welch's correction was used to compare between untreated and DS18561882-treated conditions in each cell line (*p<0.01, ** p<0.01, ***p < 0.001, ****p < 0.0001). **F-H.** Flow cytometry analysis to quantify total number of live cells in (F) MLL-AF9 (G) DF, and **H.** MOLM14 cells were analyzed for the change in their MitoSOX Median Fluorescence Intensity (MFI) by flow cytometry following treatment with either vehicle, 1 µM or 2 µM DS18561882. An unpaired t-test with Welch's correction was used to compare between untreated and DS18561882-treated conditions in each cell line (*p<0.01, ** p<0.01, ***p < 0.001, ****p < 0.0001). **H.** MOLM14 AML cells were analyzed for Annexin V staining by flow cytometry following treatment with MitoTEMPO alone or in combination 1 µM DS18561882. An unpaired t-test with Welch's correction was used to compare MitoTempo effect in the vehicle and DS18561882-treated conditions (*p<0.01, ** p<0.01).

### Supplementary Figure 6

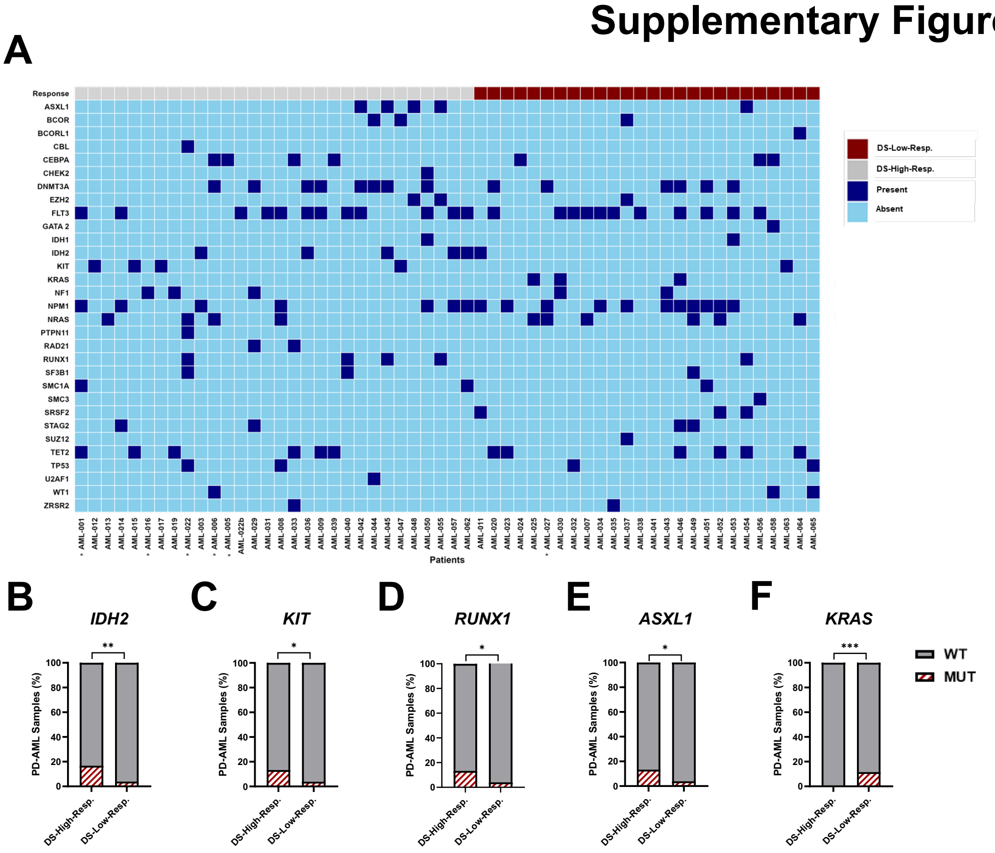


**Supplementary Figure 6**. **A.** Checkered plot showing association of mutations in 56 PD-AML samples with response to 2 μM DS18561882. **Samples denotes that analysis is based on 1 μM DS18561882.* **B-F.** Chi-square test comparison of association of (B) IDH2, (C) KIT, (D) RUNX1, (E) ASLX1 and (F) KRAS mutations with response to DS18561882. (* $p < 0.05$, ** $p < 0.01$, ****$p < 0.0001$).

### Supplementary Figure 7

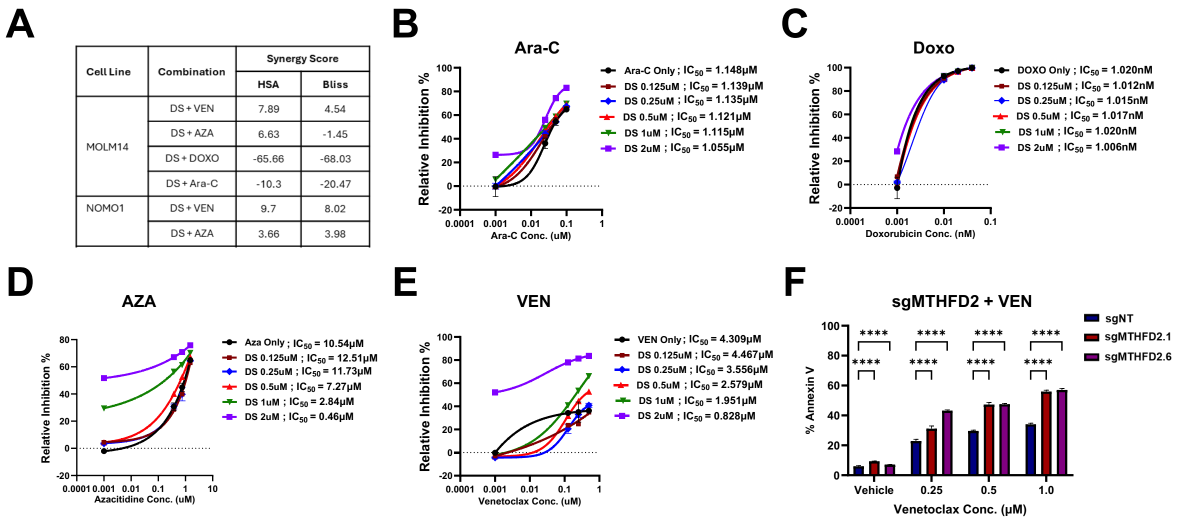


| Cell Line | Combination | Synergy Score | |
|---|---|---|---|
| | | HSA | Bliss |
| MOLM14 | DS + VEN | 7.89 | 4.54 |
| | DS + AZA | 6.63 | -1.45 |
| | DS + DOXO | -65.66 | -68.03 |
| | DS + Ara-C | -10.3 | -20.47 |
| NOMO1 | DS + VEN | 9.7 | 8.02 |
| | DS + AZA | 3.66 | 3.98 |

**Supplementary Figure 7**. **A.** Comparison of synergy scores of drug combinations in MOLM14 and NOMO1 cell lines following treatment for 96 hours. **B-E.** Comparison of cooperation between DS18561882 (DS) in combination with either (B) Cytarabine (Ara-C) and (C) Doxorubicin, (D) Azacytidine (Aza) or (E) Venetoclax (VEN) in MOLM14 human AML cells following treatment for 96 hours. **F.** MOLM14 MTHFD2 deleted (sgMTHFD2-1 and sgMTHFD2-6) and control (sgNT) AML cells were analyzed for Annexin V staining by flow cytometry following treatment with increasing concentrations of Venetoclax. An unpaired t-test with Welch's correction was used to compare the percentage of Annexin V positive cells between sgNT and each shMTHFD2 condition with or without Venetoclax treatment (****$p < 0.0001$).

#### Supplementary Figure 8

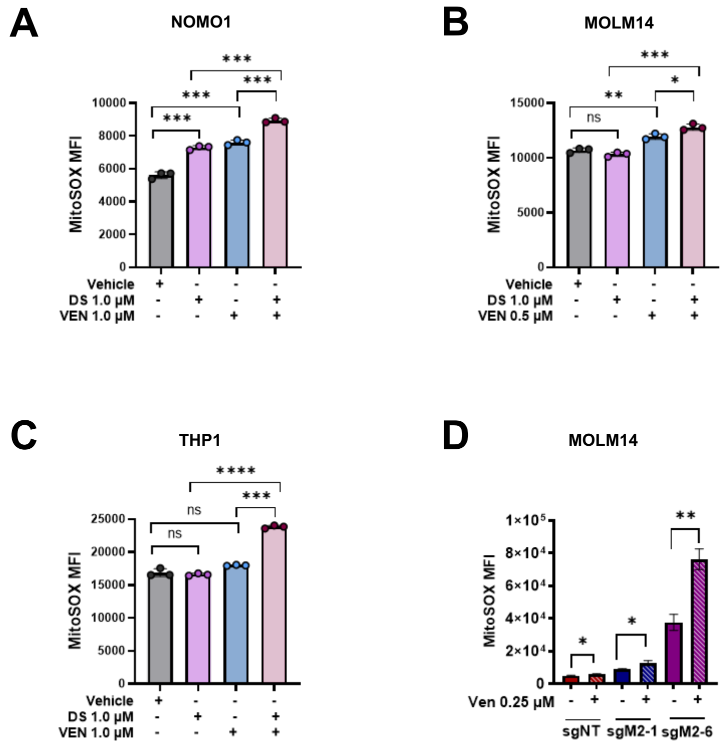


**Supplementary Figure 8. A-C.** Mitochondrial superoxide levels were assessed by MitoSOX median fluorescence intensity (MFI) by flow cytometry after treatment with vehicle or 1 μM DS18561882, with or without Venetoclax (0.5 or 1 μM) in (A) NOMO1, (B) MOLM14 and (C) THP1 Human AML cell. An unpaired t-test with Welch's correction was used to compare the difference in MitoSOX MFI of cells between vehicle or 1μM DS18561882 condition, with or without Venetoclax treatment (*p < 0.05, **p < 0.01, ***p < 0.001, ***p < 0.0001). **D.** MOLM14 cells were analyzed for change in their MitoSOX MFI by flow cytometry following treatment with either vehicle or 0.25 μM Venetoclax in MTHFD2 deleted (sgMTHFD2-1 or sgMTHFD2-6) or control (sgNT) cells. An unpaired t-test with Welch's correction was used to compare the difference in MitoSOX MFI of cells between sgNT and sgMTHFD2 conditions, with or without Venetoclax treatment (*p < 0.05, **p < 0.01).

### Supplementary Figure 9

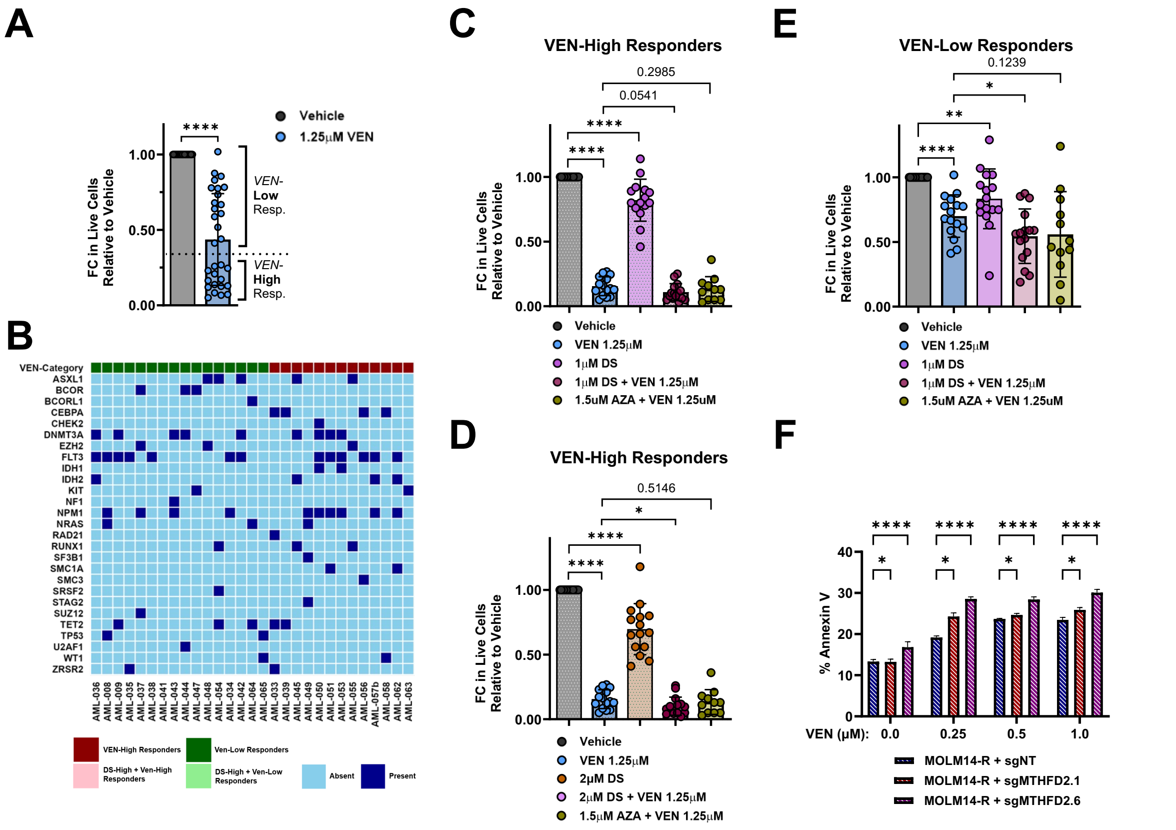


**Supplementary Figure 9. A.** Response of 31 PD-AML samples to treatment with 1.25 µM Venetoclax for 96 hours showing the Ven-High and Ven-Low repsonse category. An unpaired, nonparametric *t*-tests were used to compare vehicle and Venetoclax treated groups (****p < 0.0001). **B.** Checkered plot showing mutation profile and response of 29 PD-AML samples to treatment with 1.25 µM Venetoclax for 96hours**.** **C.** Fold change in live cells in VEN-High Responders following treatment with either 1 µM DS18561882 (DS), 1.25 µM Venetoclax (VEN), combination of 1 µM DS and 1.25 µM Venetoclax or combination of 1.25 µM VEN and 1.5 µM Azacitidine (AZA). An unpaired, nonparametric *t*-tests were used to compare vehicle and treated groups (***p < 0.0001). **D.** Fold change in live cells in VEN-High Responders following treatment with either 2 µM DS18561882 (DS), 1.25 µM Venetoclax (VEN), combination of 2 µM DS and 1.25 µM Venetoclax or combination of 1.25 µM VEN and 1.5 µM Azacitidine (AZA). An unpaired, nonparametric *t*-tests were used to compare vehicle and treated groups (* $p<0.05$, ****$p < 0.0001$). **E.** Fold change in live cells in VEN-Low Responders following treatment with either 1 μM DS18561882 (DS), 1.25 μM Venetoclax (VEN), combination of 1 μM DS and 1.25 μM Venetoclax or combination of 1.25 μM VEN and 1.5 μM Azacitidine (AZA). An unpaired, nonparametric *t*-tests were used to compare vehicle and treated groups (* $p<0.05$, **$p < 0.01$, ****$p < 0.0001$). **F.** MTHFD2 deleted (sgMTHFD2-1 and sgMTHFD2-6) and control (sgNT) MOLM14 resistant (MOLM14-R) AML cells were analyzed for Annexin V staining by flow cytometry following treatment with increasing concentrations of Venetoclax. An unpaired t-test with Welch's correction was used to compare the percentage of Annexin V positive cells between sgNT and each shMTHFD2 condition with or without Venetoclax treatment (*$p < 0.05$, ****$p < 0.0001$).

### Supplementary Table 1

| S/N | Synergy Scores | | | | | |
|---|---|---|---|---|---|---|
| | Patient ID | VEN Category | DS + Ven | | Ven + Aza | |
| | | | HSA | Bliss | HSA | Bliss |
| 1 | AML -034 | LOW | 24.08 | 24.08 | -15.9 | -18.1 |
| 2 | AML-036 | LOW | 15.33 | 7.63 | 2.61 | 0.57 |
| 3 | AML-041 | LOW | 23.83 | 6.41 | 10.51 | 0.5 |
| 4 | AML-043 | LOW | 11.2 | 9.46 | 4.89 | 4.89 |
| 5 | AML-044 | LOW | 39.1 | 34.14 | 15.41 | 12.66 |
| 6 | AML-047 | LOW | 25.29 | 20.24 | 13.35 | 8.544 |
| 7 | AML-008 | LOW | 26.58 | 16.47 | | |
| 8 | AML-009 | LOW | 45.45 | 36.53 | | |
| 9 | AML-048 | LOW | 11.45 | 9.06 | | |
| 10 | AML-037 | LOW | 5.15 | 5.15 | 13.33 | 10.82 |
| 11 | AML-042 | LOW | 2.24 | 0.61 | 5 | 2.654 |
| 12 | AML-035 | LOW | -6.49 | -6.65 | | |
| 13 | AML-038 | LOW | -0.69 | -1.49 | 3.73 | 1.81 |
| 14 | AML-039 | HIGH | 0.49 | -1.18 | 0.75 | 0.36 |
| 15 | AML-045 | HIGH | 2.39 | -0.47 | 0.58 | -0.34 |
| 16 | AML-049 | HIGH | 1.309 | 1.8 | 7.53 | 10.64 |
| 17 | AML-050 | HIGH | -1.3 | -1.3 | 1.97 | 0.84 |
| 18 | AML-033 | HIGH | 0.85 | -0.68 | | |
| 19 | AML-051 | HIGH | 4.64 | 2.32 | | |

**Supplementary Table 1.** Synergy scores of 19 PD-AML samples treated with either DS and VEN combination or VEN and AZA combination.

### Supplementary Table 2

| Name | Sequence |
|---|---|
| sgNT Fwd | CACCGATTCGTCGACGTAGGTTTCC |
| sgNT Rev | AAACGGAAACCTACGTCGACGAATC |
| sgMTHFD2-1 Fwd | CACCGCGAAGGGAGCAGCTGTGCGC |
| sgMTHFD2-1 Rev | AAACGCGCACAGCTGCTCCCTTCGC |
| sgMTHFD2-6 Fwd | CACCGACCAGGATCACACTCAGGTG |
| sgMTHFD2-6 Rev | AAACCACCTGAGTGTGATCCTGGTC |

**Supplementary Table 2.** Human *MTHFD2* and non-targeting control sgRNA oligonucleotide sequences.
