## Supplementary material for "Targeting MTHFD2 disrupts mitochondrial redox homeostasis and restores venetoclax sensitivity in acute myeloid leukemia": 4_SUPPLEMENTARY_FILE_JSSMS31626

### Materials and Methods

#### Reagents

Venetoclax and Azacitidine were purchased from Selleck Chemicals (Houston, TX, USA) and dissolved in DMSO to 10 mM stock concentrations, then stored at −80 ℃. DS18561882 (DS) was purchased from MedChemExpress (HY-130251). U-$^{13}$C-serine and 2,3,3-$^{2}$H-Serine were purchased from Cambridge Isotope Laboratories Inc., L-Serine-$^{13}$C3 was purchased from Sigma-Aldrich (St. Louis, MO, USA).

#### Cell culture and lentiviral infections

All leukemia cell lines were obtained either from the American Type Culture Collection or the German Collection of Microorganisms and Cell Cultures (DSMZ). Human cell lines were cultured in RPMI medium supplemented with 10% tetracycline-depleted fetal bovine serum and penicillin/streptomycin. In contrast, the murine cell lines MLL-AF9 were cultured in the above-described RPMI medium supplemented with 10 ng/mL murine stem cell factor (Peprotech), 6 ng/mL mIL-6 (Peprotech), and 5 ng/mL mIL-3 (Peprotech). All cell lines were authenticated by IDEXX BioAnalytics and periodically monitored in-house to ensure that they were mycoplasma-free using Hoechst staining. Control and MTHFD2-targeting sgRNAs (Supplementary Table 2) were cloned into the LRG (Lenti_sgRNA_EFS_GFP) vector, which was a gift from Christopher Vakoc (Addgene plasmid # 65656; http://n2t.net/addgene:65656; RRID: Addgene_65656)[1] while the lentiCas9-Blast was a gift from Feng Zhang (Addgene plasmid # 52962; http://n2t.net/addgene:52962; RRID: Addgene_52962). The CRISPR/Cas9 constructs, packaging, and envelope plasmids—psPAX and pMD, respectively, were transfected into HEK293TL cells using X-tremeGENE (Sigma-Aldrich) to produce high-titer lentiviral media as previously described. Human leukemia cell lines were transduced with sgRNA-expressing recombinant lentiviruses in the presence of polybrene (8 μg/mL). Stable cell lines were generated via Blasticidin selection (10 μg/mL for 72 h immediately after transduction, then maintained in culture at 10 μg/mL).

**Apoptosis assay**

Cells were washed in PBS and stained with Annexin V and Propidium Iodide (PI) according to the manufacturer's instructions (BD Biosciences). Cells were acquired and analyzed using CytoFlex (Beckman Coulter).

*Bone marrow transplant leukemia model*

All animal studies conducted were approved by the IACUC of the Washington University School of Medicine (Protocol #24-0186). The mouse strain used for this research project, C57BL/6N-$A^{tm1Brd}$ $Mthfd2^{tm1a(EUCOMM)Wtsi}$/BayMmucd, RRID: MMRRC_041551-UCD, was obtained from the Mutant Mouse Resource and Research Center (MMRRC) at the University of California at Davis, an NIH-funded strain repository, and was donated to the MMRRC by Arthur Beaudet, M.D., Baylor College of Medicine. Mice were generated at the Baylor College of Medicine as part of the Baylor College of Medicine, Sanger Institute, and MRC Harwell (BaSH) Consortium for the NIH Common Fund program for Knockout Mouse Production and Cryopreservation (1U42RR033192-01) and Knockout Mouse Phenotyping (1U54HG006348-01). Leukemia mice were generated with recombinant retroviruses expressing both MLL-AF9 and a neomycin resistance cassette as described previously[2,3]. For recombinant viral transduction, bone marrow cells recovered from leukemia mice were cultured in CEM media overnight. The next day, cells were counted, and then 500,000 cells were spin-infected with recombinant lentiviruses supplemented with polybrene (8 mg/mL) in 12-well non-adherent plates. Plates were centrifuged at 2,200 rpm for 90 minutes at 300 °C. Transduced cells were then incubated for 3 hours, after which the viral supernatant was removed, and cells were replenished with fresh CEM. For *in vivo* survival assays, sorted GFP+ cells (150,000 cells/mouse) were transplanted into sub-lethally irradiated (450 rad) syngeneic recipient mice 48 hours post-transduction. Recipient male mice were randomized into experimental groups based on similar age (8-12 weeks), weight, and vendor, and animal studies were not blinded. For western blot analysis and RNA sequencing, cells were subjected to FACS to isolate GFP+ cells following lineage depletion.

**Competitive Bone Marrow Transplant Studies**

BM cells from Mx1-Cre; Mthfd2$^{+/+}$ (wild-type), Mthfd2$^{+/-}$ (heterozygous), and Mthfd2$^{-/-}$ (homozygous knockout) mice, all expressing CD45.2. These cells were mixed at a 1:1 ratio with wild-type CD45.1$^{+}$ competitor BM and transplanted into lethally irradiated CD45.1$^{+}$ recipient mice (n=6 per genotype). Mice were treated with poly(I:C) to induce Cre-mediated deletion of Mthfd2 and subsequently monitored for donor chimerism and hematopoietic output. PB was analyzed every 5 weeks post-transplantation (up to 20 weeks) for CD45.1/CD45.2 chimerism and mature lineage distribution (myeloid, B cells, T cells, and erythroid cells).

**Patient and healthy donor samples**

Human sample collection was approved by the Washington University in Saint Louis Institutional Review Board, protocol #201011766. All human studies were conducted in accordance with the “Declaration of Helsinki” principles. Deidentified samples were cultured in RPMI-1640 supplemented with 10% FBS, 1% Penicillin/Streptomycin, 1%

GlutaMax, 1% nonessential amino acids, 1% sodium pyruvate, 2% HEPES, 0.9% β-mercaptoethanol, 100 ng/mL human stem cell factor, 10 ng/mL human FMS-like tyrosine kinase 3 ligand, 10 ng/mL human thrombopoietin, 10 ng/mL hIL-3, 20 ng/mL hIL-6, and 50 μg/mL Normocin for 4 to 5 days.

**Western blot analysis**

Total cell lysates were resolved by SDS-PAGE (4%–12% Bis-Tris gels, Thermo Fisher Scientific). Proteins were transferred to a polyvinylidene fluoride membrane, and after blocking in 1× tris-buffered saline with 0.1% Tween-20 + 5% nonfat milk for 1 h at room temperature, blots were incubated with primary antibody overnight at 4 °C. Staining with secondary antibody was performed at room temperature for 40 min and blots we developed with SuperSignal™ West Pico PLUS Chemiluminescent Substrate (Thermo scientific). The following antibodies were used: MTHFD2 (Proteintech, #10627-1-AP and Invitrogen #MA5-51284); Human FLAG-tag Cas9 (Cell Signaling, Danvers, #2209); β-Actin (Cell Signaling, #8H10B10); Tubulin (Sigma #T9206); Catalase (Cell Signaling #12980); SOD2 (Cell Signaling #13194); and MCL1 (Cell Signaling #5453). Antibodies were used and validated according to the manufacturer's instructions.

**Lentiviral transduction**

Lentiviruses were packaged in 293TL cells by co-transfection with the pPAX and pMD vectors. Cells transduced with lentiviruses expressing GFP were purified by FACS at the indicated times following transduction. 500,000 cells were transduced with recombinant pLKO.1 lentiviruses co-expressing GFP with sgRNAs from the Addgene sgRNA library[4].

**Metabolite Tracing**

Metabolomics analyses were performed at The Wistar Institute's Proteomics and Metabolomics Shared Resource. MOLM-14-cas9 cells were cultured in RPMI-1640 base media + 10% dialyzed FBS and penicillin/streptomycin, and 10μg/mL Blasticidin. Cells were transduced with recombinant lentiviruses co-expressing GFP and sgRNAs, and 24 hours later, the media was changed, and fresh media containing doxycycline (1 μg/mL) was added to induce Cas9 expression. Cells were cultured for 48 hours, after which the media was changed, and cells were then cultured in glucose-free RPMI-1640 media supplemented with 11.1 mM uniformly labeled U-13C-Glucose or serine-free RPMI-1640 with 286 mM uniformly labeled U-13C-Serine or D3- Serine for 24 hours to reach isotopic steady-state. Unlabeled controls were also used for each sgRNA condition. Polar metabolites were extracted using an ice-cold extraction solution containing 80:20 (v/v) MeOH/water. Samples were analyzed by LC-MS on a Thermo Scientific Q Exactive Plus mass spectrometer coupled to a Thermo Scientific Vanquish UHPLC System. The unlabeled sample was also analyzed by LC-MS/MS for metabolite annotation. For each analysis, 4 μL were injected per run. Samples were analyzed in a pseudorandomized order. LC separation was performed under HILIC conditions using a ZIC-pHILIC column (150x 2.1 mm, 5 μm, EMD Millipore) maintained at 45°C. Mobile phase A was 20 mM ammonium carbonate, 0.1% ammonium hydroxide, pH 9.2 and 5μM medronic acid, while mobile phase B was acetonitrile. Analytical separation was performed at 0.2 mL/min flow rate using the following gradient: 0 min, 85% B; 2 min, 85% B; 17 min, 20% B; 17.1 min, 85% B; and 26 min, 85% B. Samples were analyzed by either Full MS scans with polarity switching (all samples) or Full MS/data-dependent MS/MS scans with separate acquisitions for positive and negative polarities. Relevant MS parameters include: sheath gas, 40; auxiliary gas, 10; sweep gas, 2; auxiliary gas heater temperature, 350 °C; spray voltage, 3.5/3.2 kV for positive/negative polarities; capillary temperature, 325 °C; S-lens RF, 50. Full MS scans were acquired using a scan range of 65 to 975 m/z; 70,000 resolution; automated gain control (AGC) target of 1E6; and maximum injection time (IT) of 100 ms. Data-dependent MS/MS was performed on the 10 most abundant ions; 17,500 resolution; AGC target of 5E4, maximum IT of 50 ms, isolation width of 1.0 m/z, and stepped normalized collision energy of 20, 40, 60. Raw data were processed using Compound Discoverer (Thermo Scientific) with separate analyses for positive and negative polarity. Metabolites were identified by matching accurate mass and retention time to standards or querying MS/MS spectrum against the mzCloud spectral database (full match, score > 50; mzCloud.org). Identifications were transferred to labeled samples in isotope-tracing experiments, accounting for all possible $^{13}C$ isotopologues. Metabolite levels were expressed as MS signal, i.e. integrated peak areas using Full MS data. Isotope tracing data were corrected for natural $^{13}C$ abundance and isotope tracer purity. Metabolite levels were further normalized to total protein recovered from the polar metabolite extraction pellet.

Redox Staining

Human AML cells expressing either sgNT, MTHFD2.1, or MTHFD2.6 were washed and incubated with PBS + 5 mmol/L of MitoSOX or CellROX DeepRed reagent (Life Technologies; catalog no.: C10422) at $37^0$C for 10minutes. After incubation, cells were washed twice with PBS and analyzed using a CytoFlex flow cytometer (Beckman Coulter).

**Nucleotide supplementation**

Human AML cells expressing MTHFD2-targeting sgRNAs or control sgRNA were seeded at a concentration of 500,000 cells/mL in 12-well plates and treated every other day with 25 µm of 2'-deoxyadenosine (Sigma Aldrich, D8668) and 2'-deoxyguanosine (Sigma Aldrich, D0901) for up to 15 days and then analyzed by flow cytometry.

**Quantification and Statistical Analysis**

Data presented for the *in vitro* experiments using mouse or human AML cell lines are representative of a single experiment and are expressed as mean ± SD from 3 technical replicates; each experiment was repeated at least 3 times. The *in vivo* experiments utilizing a mouse AML cell line are representative of a single experiment and are expressed as mean ± SD of at least 3 biological replicates, and each experiment was repeated at least twice. For PD-AML studies, the presented data represents the mean ± SD of three wells from a single experiment. Comparisons between two experimental groups were performed using Student's t-test (unpaired 2-tailed) unless otherwise indicated, and significance was determined using the following p-values: * $p < 0.05$, ** $p < 0.01$, *** $p < 0.001$, **** $p < 0.0001$. P values from all survival curves are analyzed using the Log-rank (Mantel-Cox) test to compare two survival curves. Statistical tests were performed using GraphPad Prism 10 software.
